# Dynamics of the compartmentalized *Streptomyces* chromosome during metabolic differentiation

**DOI:** 10.1101/2020.12.09.415976

**Authors:** Virginia Lioy, Jean-Noël Lorenzi, Soumaya Najah, Thibault Poinsignon, Hervé Leh, Corinne Saulnier, Bertrand Aigle, Sylvie Lautru, Annabelle Thibessard, Olivier Lespinet, Pierre Leblond, Yan Jaszczyszyn, Kevin Gorrichon, Nelle Varoquaux, Ivan Junier, Frédéric Boccard, Jean-Luc Pernodet, Stéphanie Bury-Moné

## Abstract

*Streptomyces* are among the most prolific bacterial producers of specialized metabolites, including antibiotics. The linear chromosome is partitioned into a central region harboring core genes and two extremities enriched in specialized metabolite biosynthetic gene clusters (SMBGCs). The molecular mechanisms governing structure and function of these compartmentalized genomes remain mostly unknown. Here we show that in exponential phase, chromosome structure correlates with genetic compartmentalization: conserved, large and highly transcribed genes form boundaries that segment the central part of the chromosome into domains, whereas the terminal ends are transcriptionally, largely quiescent compartments with different structural features. Onset of metabolic differentiation is accompanied by remodeling of chromosome architecture from an ‘open’ to a rather ‘closed’ conformation, in which the SMBGCs are expressed forming new boundaries. Altogether, our results reveal that *S. ambofaciens’* linear chromosome is partitioned into structurally distinct entities, indicating a link between chromosome folding, gene expression and genome evolution.

## Introductory paragraph

Bacteria of the genus *Streptomyces* are amongst the most prolific producers of specialized metabolites with applications in medicine, agriculture and the food industry^1^. The biosynthesis of these specialized metabolites (*e.g*. antibiotics and pigments) generally occurs after ‘metabolic differentiation’, a physiological transition from primary to specialized metabolism^2^,^3^. This transition coincides with morphological differentiation^4^ (*e.g*. formation of the secondary multinucleated mycelium, sporulation) that occurs late in the growth phase. These metabolic and morphological changes are controlled by highly interconnected regulatory mechanisms that have been studied mostly at the transcriptional, translational and post-translational levels^4–7^. However, a clear link between global chromosome organization and metabolic differentiation has yet to be established.

*Streptomyces* possess an unusual linear chromosome. Moreover, its extremities, or ‘terminal arms’, contain terminal inverted repeats (TIRs) capped by telomere-like sequences. In addition, the chromosome is large (6-15 Mb) with an extreme GC content (circa 72%). Finally, the *Streptomyces* chromosome presents a remarkable genetic compartmentalization, with a distinguishable central region and the terminal arms. The central region primarily contains core genes, common to all *Streptomyces* species, these being often essential^8–13^. The terminal arms are mainly composed of conditionally adaptive genes, in particular, enriched in specialized metabolite biosynthetic gene clusters (SMBGCs)^14^. In addition, they are prone to large DNA rearrangements and frequent recombination^12^,^15^,^16^,^17^. Importantly, many SMGBCs appear silent or poorly expressed under laboratory conditions, giving rise to the concept of ‘cryptic’ SMBGCs. The basis of the regulation of these clusters in relation to the overall genome dynamics remains to be explored.

The development of chromosome conformation capture (3C) methods coupled to deep sequencing provided novel concepts in bacterial chromosome biology^18,19^. Notably pioneer studies revealed that bacterial chromosomes present a high degree of 3D-organization in macrodomains and/or ‘chromosome interacting domains’ (CIDs), mediated by multiple structural factors, including transcription and replication^20–27^. Here, we explore the dynamics of the *Streptomyces* chromosome during metabolic differentiation. By combining multi-omic approaches, we show that the dynamics of gene expression correlate with the folding of the linear chromosome of *Streptomyces ambofaciens* ATCC 23877 into transcriptionally active and silent compartments. Moreover, metabolic differentiation is accompanied by a huge remodeling of chromosome architecture from an ‘open’ to a rather ‘closed’ conformation, SMBGCs forming new boundaries. Altogether, our results highlight a link between chromosome folding, gene expression and genome evolution.

## Results

### Compartmentalization of the *Streptomyces ambofaciens* genome highlighted by comparative genomics and functional annotation

The genetic compartmentalization of the *S. ambofaciens* ATCC 23877 genome was previously reported^12^. However, here we took advantage of the numerous *Streptomyces* available sequences to update the cartography of the genome. We also used gene persistence^28^ as a new indicator to describe the organization of *Streptomyces* genome. Gene persistence reflects the tendency for genes to be conserved in a large number of genomes. As shown in other bacteria, gene persistence is associated with gene essentiality and similar expression levels^28–30^. Here, we calculated a gene persistence index by determining the frequency of a given gene in a panel of 125 complete *Streptomyces* genomes (**Supplementary Table 1**)^31^. The highest level of persistence associated with the best reciprocal matches between genes of all these genomes defines the core-genome^31^. In addition, we mapped the genes encoding the ‘actinobacterial signature’ previously identified^32^ as those coding sequences (CDSs) that are nearly universal among actinobacteria. In contrast, we searched the *S. ambofaciens* ATCC 23877 genome for variable regions, potentially acquired by horizontal gene transfer. For this purpose, we identified unique genes as well as genomic islands (GIs, **Supplementary Table 2, Supplementary Fig. 1**). These are defined as DNA sequences that are inserted in regions of synteny (i.e. same orthologous gene order in closely related genomes, see Methods). Thus genes from the core, the actinobacterial signature and/or presenting a high level of persistence are enriched in the central region, whereas GIs and unique CDSs are enriched in chromosome extremities. The results clearly highlight the compartmentalization of the *S. ambofaciens* genome (**Fig.1**, **Supplementary Table 3**). Synteny gradually disappears in terminal arms (**Fig.1**), as previously reported^12^. This makes it difficult to have an operational delineation of the limits of these arms. However, since the first and last ribosomal operons approximately mark the limits of a central region beyond which the synteny level falls (**Fig.1**), these operons were used as limits for definition of the left and right extremities in this study.

**Figure 1:**
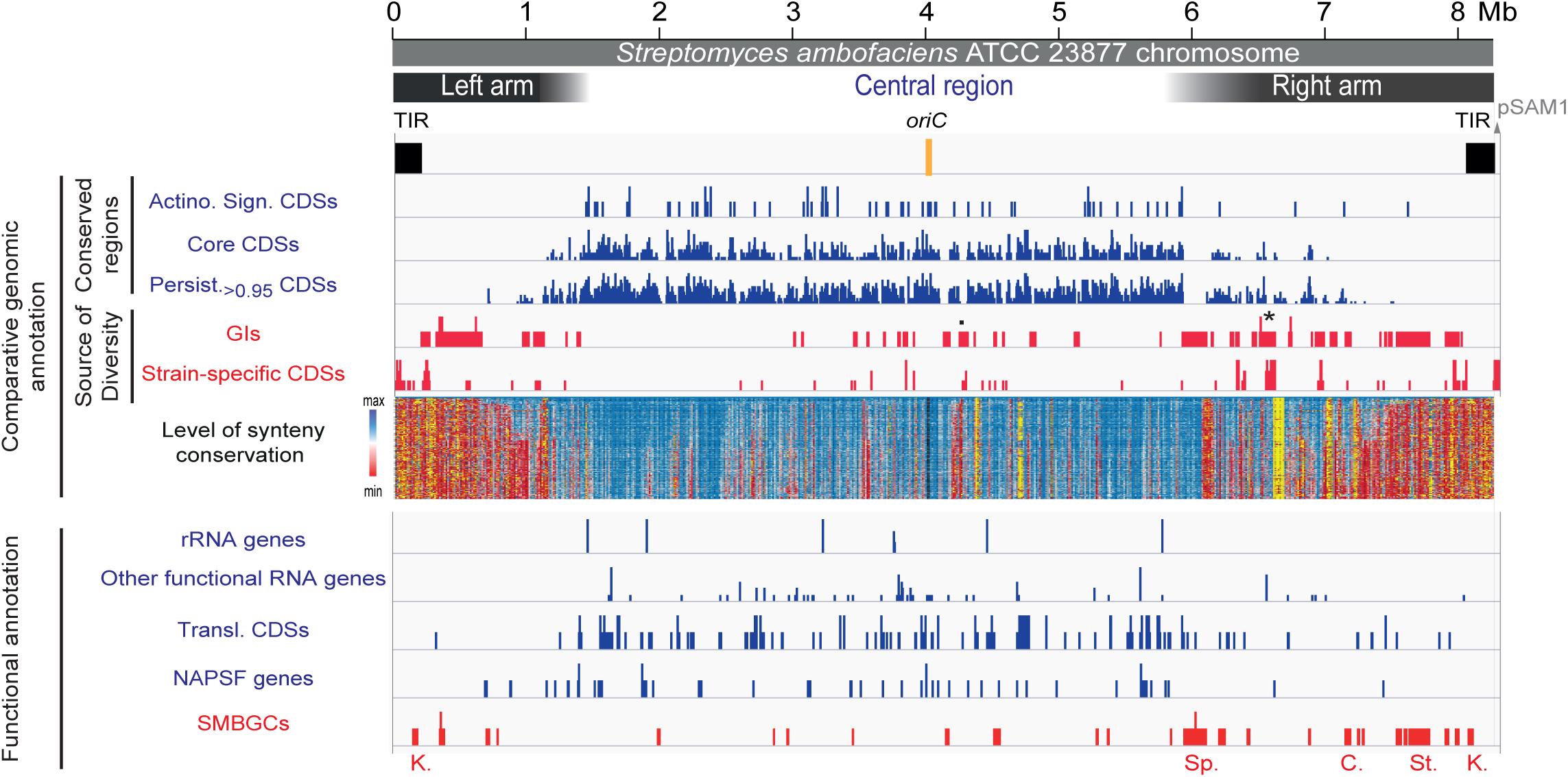
The genetic compartmentalization of *S. ambofaciens* linear chromosome. The terminal inverted repeats (TIRs, 200 kb) and the origin of replication (*oriC*) are indicated in black. The distribution of gene presenting features of interest are represented in blue or red when enriched in the central region or within the extremities (defined by the first and last rDNA operons), respectively (see **Supplementary Table 3** for a statistical analysis of the data). The dot and asterisk indicate the position of pSAM2 and of a complete prophage, respectively. The position of the SMBGC encoding the biosynthesis of all known antibacterial compounds of *S. ambofaciens* ATCC 23877 is also indicated. The level of synteny along the chromosome of *S. ambofaciens* ATCC 23877 is represented as a heat map of GOC (Gene Order Conservation) scores using a sliding window (8 CDSs with 1 CDS steps). Each line corresponds to the GOC profile of the reference against another species. The species are organized from the phylogenetically closest to the furthest compared to *S. ambofaciens*. The vertical black dotted line represents the location of the *dnaA* gene delineating the two replichores. “NA” (in yellow) indicates the absence of orthologs in the corresponding window. *S. ambofaciens* ATCC 23877 also harbors a circular pSAM1 plasmid (≈ 89 kbp). Abbreviations: ‘Actino. Sign. CDSs’ (coding sequences of the actinobacterial signature); BGC (biosynthetic gene cluster); ‘C.’ (congocidine BGC); ‘Chr.’ (whole chromosome); GIs (genes belonging to genomic islands); ‘K.’ (kinamycin BGC); ‘Persist.>0.95 CDSs’ (coding sequences of *S. ambofaciens* ATCC 23877 presenting a gene persistence superior to 95 % in 124 other *Streptomyces* genomes); ‘Transl. CDSs’ (genes encoding functions involved in translation process and/or RNA stability); NAPSFs (nucleoid associated proteins and structural factors); SMBGCs (specialized metabolite BGCs); ‘Sp.’ (spiramycin BGC); ‘St.’ (stambomycin BGC); ‘rRNA’ (ribosomal RNA).

Functional annotation also highlighted a bias of gene distribution along the chromosome (**Fig.1**, **Supplementary Table 3**). As a proxy for major metabolic processes, we used the functional RNA genes (encoding rRNA, tRNA, tmRNA, RNAse-P RNA, SRP-RNA) and genes encoding functions related to translation and/or RNA stability. We also mapped the genes encoding nucleoid-associated proteins (NAPs) and chromatin structural factors, further shorten ‘NAPSFs’, that play a central role in the dynamic organization of the bacterial chromosome and are enriched in the central region (**Fig.1**, **Supplementary Table 3**). Finally, we used the antiSMASH secondary metabolite genome mining pipeline (version 5.1.0)^33^ to identify putative SMBGCs (**Supplementary Table 4**). These genes are preferentially located in the terminals arms and almost half of them are present within GIs (**Fig.1**, **Supplementary Table 3**). Accordingly, GIs are 3.5-fold enriched in SMBGC genes (*p* value < 2.2.10^−6^, Fisher’s exact test for count data). These results illustrate that the high variability of *Streptomyces* extremities relates in particular to functions involved in metabolic differentiation.

Together, these analyses confirm the strong genetic compartmentalization of the *S. ambofaciens* chromosome. In this context, we then explored the extent to which this genetic organization correlates with gene expression and chromosome architecture.

### Transcription of the *S. ambofaciens* genome is strongly compartmentalized

To focus specifically on transcriptome dynamics during metabolic differentiation, while limiting cellular physiological heterogeneity in the colony, we took advantage of the fact that *S. ambofaciens*, like many *Streptomyces^2^*, does not sporulate in liquid medium. We chose MP5 and YEME liquid media (**Table 1**) in which *S. ambofaciens* grows in a rather dispersed manner that makes the cells also more accessible to further 3C-treatments. Moreover, for *S. ambofaciens*, MP5 medium was previously reported to be suited for the production of the antibiotics spiramycin^34^ and congocidine^35^ by *S. ambofaciens* while there is only limited antibiotic production in YEME medium (**Extended Data Fig.1.A&B**). The results showed that the transcriptomes were rather similar in exponential phase in both media, while commitment to specialized metabolism (C4, C5 and C7 conditions) was accompanied by major transcriptional changes that form a distinct cluster (**Fig.2.A&B, Extended Data Fig.1.C&D**). Notably, at the earliest time points in the growth period in both media, the terminal regions were rather poorly expressed, whereas transcription gradually increased toward terminal ends over the growth period (**Fig.2.B**), as previously reported in *S. coelicolor*^10^,^36^. We observed that the *S. ambofaciens* genome has a transcriptional landscape with approximately 90 % of the genes significantly expressed (‘CAT_1’ level or more) and/or regulated (DESeq2 statistical analysis) in at least one condition (**Extended Data Fig.2.A&B**, **Supplementary Table 5**).

**Figure 2:**
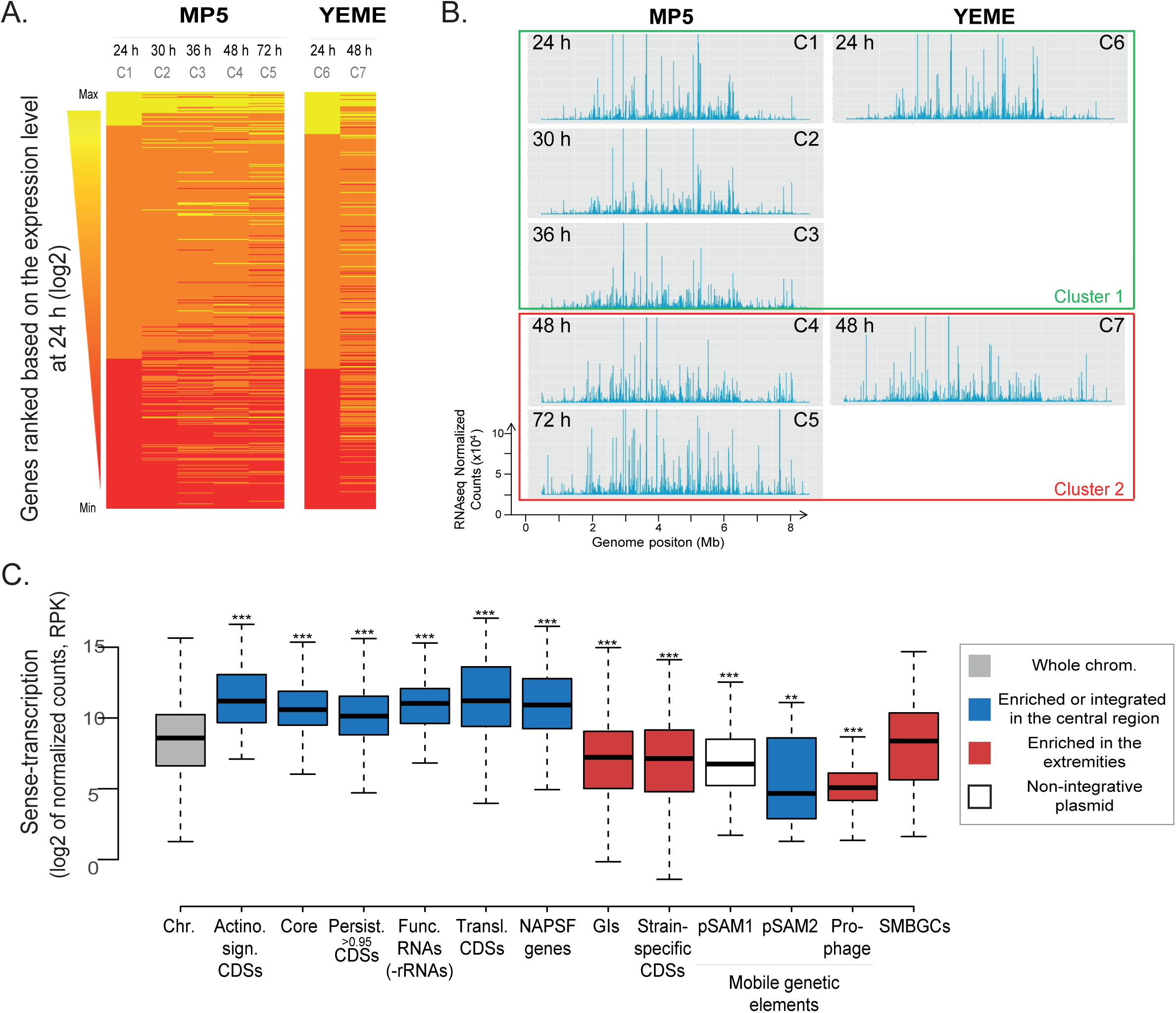
Transcriptome dynamics depending on genome features and metabolic differentiation. **A. Transcriptome changes over growth.** Heatmaps of the normalized number of reads per gene, ranked by their expression level at 24 h. **B. Transcription along the chromosome over growth.** The normalized counts (DESeq2 analysis) were mapped on *S. ambofaciens* chromosome. The two clusters to which the different conditions belong were obtained by hierarchical classification (**Extended Data Fig.1C**). Whereas the cluster 1 corresponds to exponential phases (no ATB production), the cluster 2 gathers conditions associated to the activation of SMBGC encoding at least two known antibacterial activities (**Supplementary Fig.2**). ‘C1’ to ‘C7’ refer to the name of the studied conditions: C1 to C5, cells grown in MP5 during 24 h, 30 h, 36 h, 48 h and 72 h, respectively; C6 and C7: cells grown in YEME during 24h and 48h, respectively (see **Table 1,** for details). **C. Boxplot presenting the mean level of gene expression depending on features of interest.** The distribution of the number of counts (normalized by DESeq and on gene size) as the mean of all studied conditions is presented depending on the genome features of interest (see the legend of **Figure 1** for abbreviations). For clarity, outliers were excluded from this graphical representation (but were taken into account for the numerical exploitation of the data). The significant differences by Wilcoxon rank sum test with continuity correction compared to the chromosome (‘Chr.’) are indicated by asterisks (**, *p*< 0.001, ***, *p*< 0.001). Other abbreviation: ‘Func. RNA (-rRNA)’ (Functional RNA excluding rRNA)

**Table 1:** Growth conditions used to performed -omics analyses

| Conditions | Media | Brief description | Time (h) | Antibacterial activity* |
| --- | --- | --- | --- | --- |
| **C1** | **MP5** | Liquid medium based on yeast extract containing glycerol (3.6 %), NaNO<sub>3</sub> (1 g/l), MOPS - optimized for congocidine and spiramycin production | 24 | - |
| **C2** |  |  | 30 | - |
| **C3** |  |  | 36 | - |
| **C4** |  |  | 48 | ++ |
| **C5** |  |  | 72 | +++ |
| **C6** | **YEME** | Liquid medium based on yeast extract containing saccharose (10.3 %), glucose (1 %), and malt extract | 24 | - |
| **C7** |  |  | 48 | + |

\*Size of the inhibition halo: “-” (no detectable inhibition), “+” (< 20 mm), “++” ([20; 30] mm), “+++” (> 30 mm) – See **Extended Data Fig.1B**.

As shown by scanning the genome with respect to gene expression levels, we found a marked bias in the localization of poorly *versus* highly expressed genes along the genome as well as a positive correlation between gene persistence and expression (**Fig. 2C, Extended Data Fig.2.C & D**). The 332 chromosomal genes expressed at very high level in all conditions are mainly enriched in the central region of the genome (**Supplementary Table 3, Fig.3D**), whereas most (> 60 %) of the SMBGCs, GIs and mobile genetic elements (pSAM1, pSAM2 and a prophage) are silent or poorly expressed in at least one condition. Up to 19 and 23 % of SMBGC and GI genes, respectively, were found in this category in all tested conditions (**Extended Data Fig.2.B**). However, compared to other poorly conserved genes, SMBGCs have a higher proportion of highly induced genes, from ‘CAT_0’ to the highest (‘CAT_3’ or ‘CAT_4’) categories (**Extended data Fig.2.C**).

**Figure 3:**
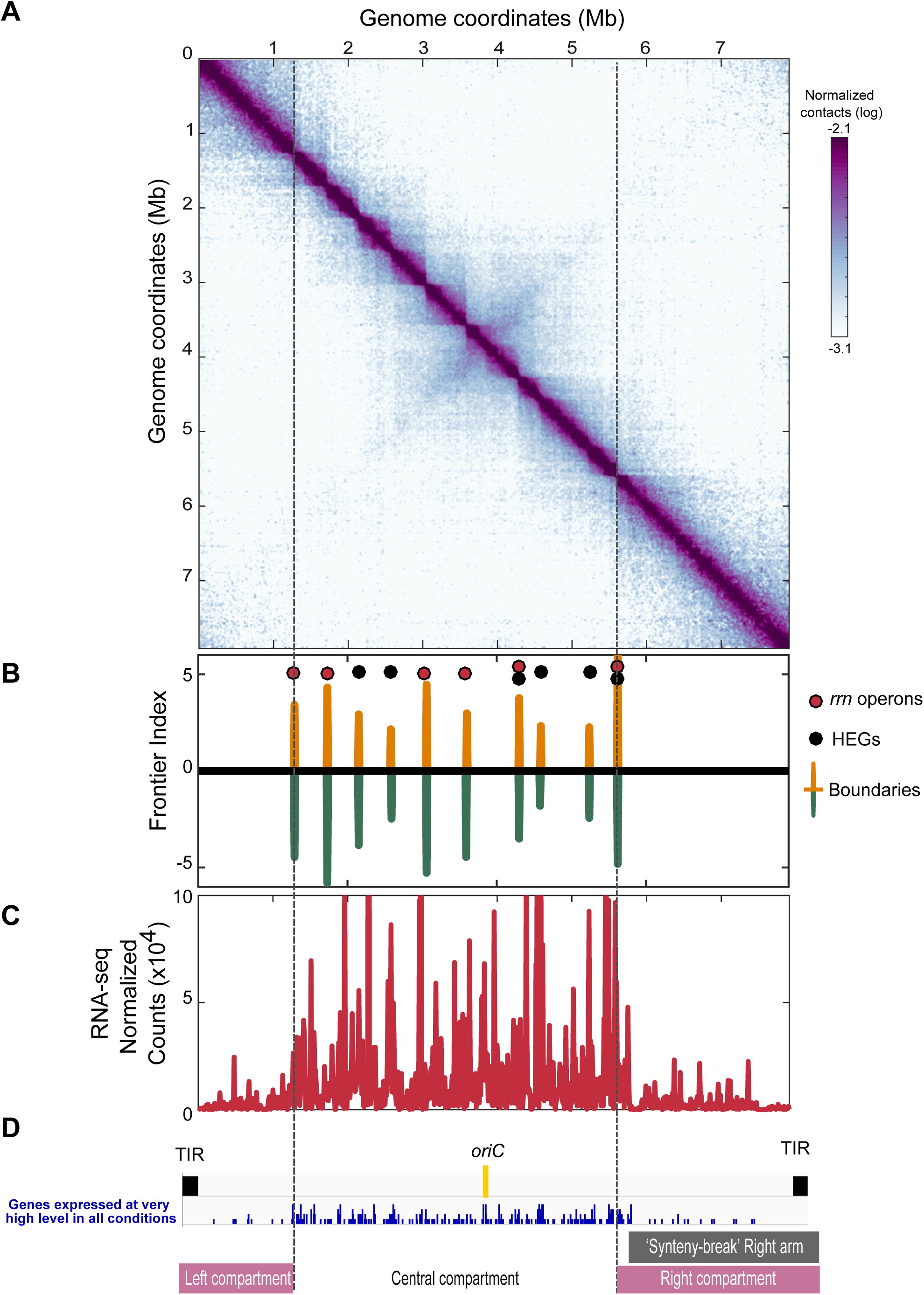
Spatial organization of *S. ambofaciens* chromosome and transcriptome in absence of metabolic differentiation. 3C-seq and RNA-seq were performed on *S. ambofaciens* grown in exponential phase in YEME medium (24 h, C6 condition). The normalized contact map was obtained from asynchronous populations. The *x* and *y* axes represent the genomic coordinates of the reference genome. To simplify the analyses, TIRs were removed from the reference genome. In **panel A**, the colour scale reflects the frequency of the contacts between genome loci, from white (rare contacts) to dark purple (frequent contacts). In **panel B**, the frontier index analysis is able to detect, for a given genome 10 kb bin, the change in the contact bias with their neighbouring bins. Thus, a boundary is defined as any bin in which there is a change in the right bias of contacts towards the left bias (± 2 bin, green and orange peaks, respectively). Red and black circles indicate the position of rDNAs and highly expressed genes (HEGs), respectively. In **panel C**, the normalized counts (DESeq2 analysis) measured in cells grown in the same condition were mapped on *S. ambofaciens* chromosome and binned in 10 kb. The **panel D** highlights genomic and transcriptional features of interest. The right and left compartment were defined owing the outer boundaries of the central compartment, which correspond to the first and last rDNA operon position. The ‘synteny-break’ right arm corresponds to the beginning of the spiramycin BGC. The replicate of this experiment is presented in the **Supplementary Fig. 3.A**.

Interestingly, as previously described in *S. coelicolor^36^*, we observed significant antisense-transcription throughout the genome. Considering the absolute number of transcripts, this antisense-transcription was positively correlated to the level of sense-transcription (**Extended Data Fig.2.E**), suggesting that spurious antisense-transcription can arise from highly transcribed regions. To investigate the relative importance of this antisense-transcription, we defined the ‘antisense index’ as the level of antisense-transcription over the total (sense plus antisense) transcription. Interestingly, in exponential phase, the antisense index was particularly high for GIs, mobile genetic elements and SMBGCs and very low in conserved regions enriched in the central region (except for pSAM2) (**Extended Data Fig.2.F**). During metabolic differentiation the antisense indices of GIs, pSAM1, prophage and especially SMBGCs tended to decrease (**Extended Data Fig.2.F**). This observation suggests that antisense-transcription could be a consequence of or directly involved in the regulation of gene expression concerning metabolic differentiation.

Together, these results indicate that the genetic compartmentalization of the *Streptomyces* genome clearly correlates with a compartmentalization of transcription, both sense and antisense.

### Compartmentalized architecture of the *Streptomyces ambofaciens* chromosome in exponential phase

We next asked the question, is a compartmentalized transcriptome correlated with a specific chromosome folding. We first performed 3C-seq on cells harvested in the exponential growth phase (**Fig.3**, **Fig.4.A**). The contact map displayed a main diagonal reflecting the frequency of contacts, which extends up to 1 Mb (**Extended Fig.3.A**). This diagonal contains loci acting as boundaries delimiting segments (visualized as squares, **Fig.3.A**, **Fig.4.A**) reminiscent of CIDs in other bacteria^20–22^. To define these domains precisely in *S. ambofaciens*, we computed the ‘frontier index’, a domain boundary indicator built from a multiscale analysis of the contact map^37^. Briefly, two indices are computed, reflecting the intensity of the loss of contact frequencies when going downwards or upwards, respectively, to each genome position (**Extended Fig.3.B**). In this context, a boundary is defined by a significant change in both the downstream (green peaks, **Fig.3.B**) and upstream (orange peaks, **Fig.3.B**) directions (see method section for details). In exponential phase in YEME medium (non-optimal for antibiotic production, condition C6), we found 10 boundaries that defined the central region, delimited by the first and last rDNA operons, 9 domains ranging in size from 240 kbp to 700 kbp (**Fig.3.B**). These central domains resembled regular CIDs^20^,^23^ both in size and in the presumably nature of its formation (see below). The central boundaries are also conserved in exponential phase in MP5 medium, which is optimized for antibiotic production (**Fig.4.A**). Only four additional boundaries are observed in this medium (**Fig.4.A**). All rDNA operons coincide with the sharpest boundaries (**Fig.3.B, Fig.4.A**). The genes surrounding these operons, as well as those of other boundaries, are enriched in consecutive genes expressed at very high level (**Fig.4.B, Supplementary Table 5**), as previously reported in other bacterial models^20^,^23^. Interestingly, in exponential phase, we report a clear correlation between the level of gene conservation and the formation of boundaries (**Fig.4.C**). Moreover, the transcription of the genes present within boundaries tends to be oriented in the direction of continuous replication (odds ratio 1.8, *p* value 1.4.10^−8^, Fisher’s exact test for count data), with a very low antisense index (**Extended Data Fig.3.C**).

**Figure 4:**
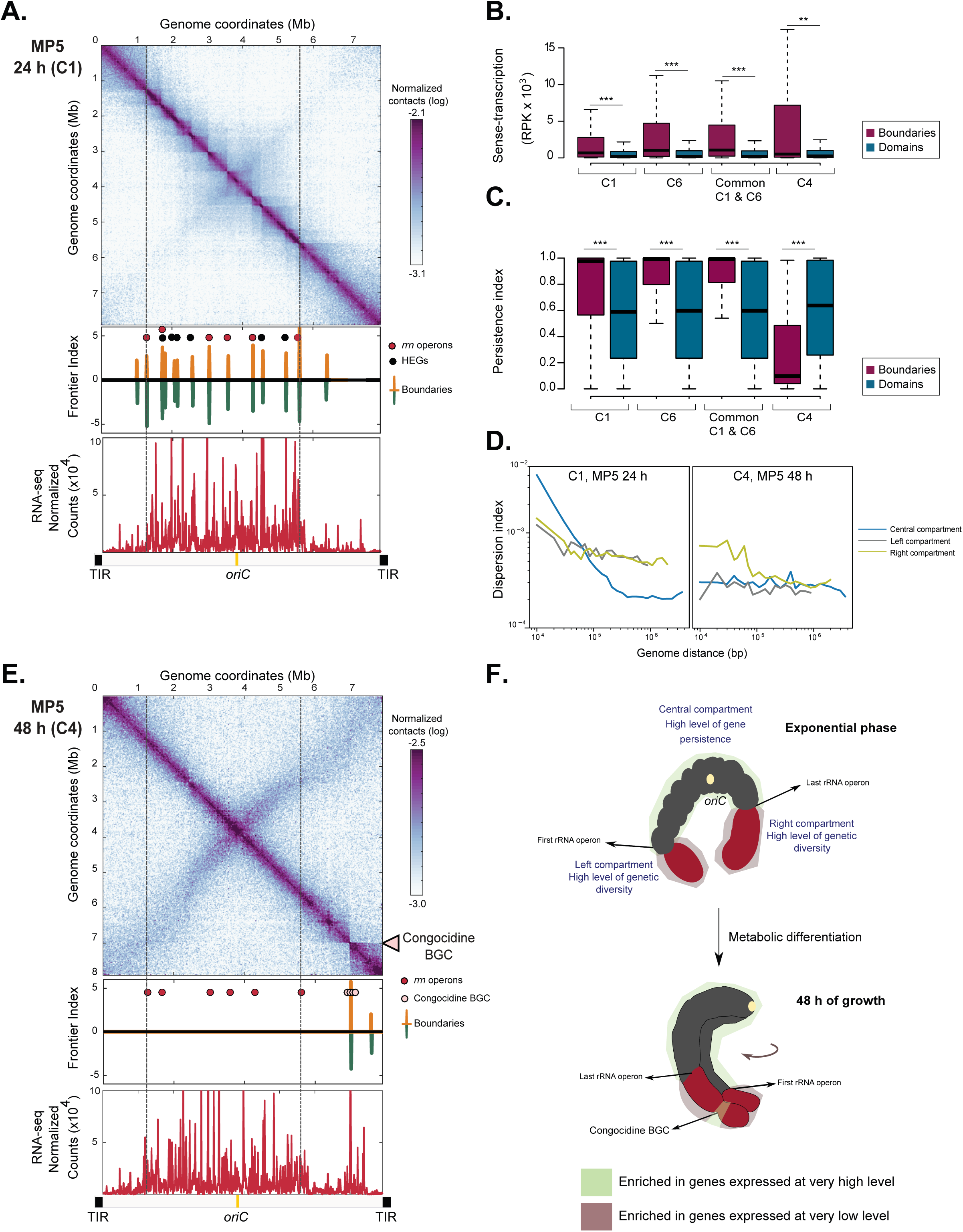
Chromosome remodeling during metabolic differentiation. **A. 3C-seq and RNA-seq performed after 24 h in MP5 growth medium** (C1 condition). Same legend than **Figure 3**. The replicate of this experiment is presented in the **Supplementary Fig. 3.B**. **B. Level of gene expression depending on gene location within boundaries or domains.** The box-plot presents the sense-transcription (normalized number of read per Kb) depending on gene location in exponential phase either in MP5 (C1) or in YEME (C6), or after 48 h of growth in MP5 medium (C4). Boundaries in common between MP5 24 h and YEME 24 h conditions (‘Common C1 & C6’) were also analyzed separately. The significant differences by Wilcoxon rank sum test with continuity correction (boundaries versus domains) are indicated by asterisks (**, *p* < 0.01; ***, *p* < 0.001). **C. Level of gene persistence depending on gene location within boundaries or domains.** The box-plot presents the persistence index depending on gene location in exponential phase (C1: MP5; C6: YEME) or after 48 h of growth in MP5 medium (C4). Boundaries in common between MP5 24 h and YEME 24 h conditions (‘Common C1 & C6’) were also analyzed separately. The significant differences by Wilcoxon rank sum test with continuity correction (boundaries versus domains) are indicated by asterisks (***, *p* < 0.001). **D. Dispersion index of the left, central and right compartments plotted as a function of genomic distance.** The index of dispersion (*Var*(*s*)/*P*(*s*)) reflects the range of variations relative to the mean value as a function of the genome distance for the contact maps obtained for cells grown 24 h and 48 h in MP5 medium. Long-range DNA contacts within the terminal compartments (> 100 kbp) are more variable than within the central compartment. **E. 3C-seq and RNA-seq performed after 48 h in MP5 growth medium** (C4 condition). Same legend than **Figure 3**. The arrow indicates the position of the congocidine biosynthetic gene cluster (BGC). The replicate of this experiment is presented in the **Supplementary Fig.3.C**. **F. Schematic representation of genome dynamics during metabolic differentiation.** The *Streptomyces* linear chromosome is represented as dark gray (central compartment) and dark red sticks (left and right compartments). The origin of replication is indicated as a light yellow circle (*oriC*). In exponential phase, the chromosome of *Streptomyces* is organized into one transcriptionally active (light green) central compartment and two rather silent compartments (pink grey). The transcriptionally active compartment is enriched in core genes and is segmented in multiple domains, flanked by long and highly expressed conserved genes. The low frequency of contacts between the two arms is represented as an open conformation of the chromosome. After 48 h of growth, the transcription of the congocidine cluster is accompanied by a global remodeling of the chromosome. The increase in the frequency of contacts between the two chromosomal regions around the origin is represented as a closed conformation. Interestingly, a slight tilt on the secondary diagonal towards the ends of the chromosome shows an asymmetry between contacts within the terminal ends that was not schematized in the figure. In addition, since TIRs cannot be distinguished at the sequence level, they were neither considered in this analysis nor represented in this figure. However they may be located close to each other^74^, although this remains controversial^75^.

Genome extremities form two large terminal compartments of 1.46 and 2.46 Mbp, respectively, with different structural features (**Fig.3, Fig.4.A** and **Extended Data Fig.3B**). Interestingly, within the terminal arms, the low transcriptional activity in exponential phase in YEME medium correlates well with the absence of boundaries. Of note, the boundaries present in each terminal domain in MP5 medium (one of which is composed of an active prophage) do not contain highly expressed genes and do not divide the terminal compartment into two parts (**Fig.4.A**). To compare the dynamics of contacts within regions, we thus calculated a dispersion index, reflecting the variability of the 3C-seq signal in each region (**Fig.4.D**, **Extended Data Fig.3.E)**. Despite that the plots for the probability of contacts as a function of genomic distance are similar for both the terminal and central regions (**Extended Data Fig.3.D**), the terminal ends present a higher dispersion index than the central compartment in both media for contacts longer than 100 kbp (**Fig.4.D**, **Extended Data Fig.3.E)**. This indicates that contacts between loci in the terminal regions are more variable than in the central compartment.

Thus, at the conformational level, the chromosome of exponentially growing cells of *S. ambofaciens* is partitioned into three compartments: a central compartment, actively transcribed and structured in multiple domains formed by the high-level expression of persistent genes and two large terminal compartments rich in GIs and SMBCGs and rather transcriptionally silent in which large-scale contacts are highly variable. Since in exponential phase, the boundaries are very similar in YEME and MP5 media and correspond mostly to persistent CDSs (**Fig.4.C**) or rDNA, their orthologous sequences may also structure the spatial organization of the chromosome in other *Streptomyces* species at a similar growth phase.

### Chromosome remodeling during metabolic differentiation

As shown above, during metabolic differentiation we observed that the level of transcription gradually increased within the terminal ends (**Fig.2**). To determine whether transcription dynamics influence the 3D-organization of the chromosome, we performed 3C-seq on *Streptomyces* cells during metabolic differentiation in MP5 medium (48 h, condition C4) (**Fig.4.E**). The frontier index revealed changes in the boundaries along the primary diagonal; the boundaries identified in exponential phase were not detected anymore after 48 h in the growth medium (**Fig.4.E**). By contrast, the appearance of a very sharp boundary within the right terminal region that divides this arm into two domains of 1,350 kpb and 1,000 kpb, correlates very well with the increased level of transcription of the congocidine biosynthetic gene cluster (BGC, **Fig.4.B & E**). In a replicate experiment, we observed that the expression of another SMBGC, encoding the biosynthesis of a siderophore, could generate the formation of an additional boundary (**Supplementary Fig.3.C**). This illustrates some variability in the expression of SMBGCs during metabolic differentiation. Together, these results highlight the correlation between SMBGC expression and the formation of boundaries after 48 h. Accordingly, the level of conservation of genes present within the boundaries switches from highly persistent in exponential phase to poorly conserved during metabolic differentiation (**Fig.4.C**). In addition, a second boundary detected within the right compartment is located in the stambomycin BGC that is not or is poorly expressed in this condition. Interestingly, accompanying the formation of boundaries and the increase of transcription in the terminal compartments, the variability in the long-range contacts within the terminal compartments is comparable to that of the central compartment (**Fig 4.D**).

The contact map also revealed the appearance of a secondary diagonal after 48 h of growth, indicating an increase in the frequency of contacts between regions along the entire arms of the chromosome. This suggests that late in the cell cycle and in the absence of a central compartment segmented into multiple domains, the two chromosome arms are closer to each other (**Fig.4.E** & **F**). Remarkably, the second diagonal is slightly tilted when it moves away from the origin, representing an asymmetry in the contacts between loci within the terminal compartments.

In summary, our results indicate that the central and transcriptionally active compartments in exponential phase present no boundaries after 48 h of growth, whereas the terminal domains are locally remodeled concomitantly with the expression of SMBGCs and with a decrease in the variability of the long-range contacts. In addition, inter-arm contacts become more frequent. These results indicate that metabolic differentiation is accompanied by major remodeling of the 3D-architecture of the chromosome, both at the local and global levels (**Fig.4.F**).

## Discussion

In this study, we demonstrated that in *Streptomyces*, compartmentalization of gene organization, transcription and architecture are correlated. We explore the transcriptional landscape of *S. ambofaciens* ATCC 23877 during metabolic differentiation: its large genome is highly dynamic, most of the genes (≈ 90 %) being significantly expressed in at least one condition. This situation is very similar to *Bacillus subtilis*, another Gram-positive bacteria from soil^38^. Interestingly, the SMBGCs (*e.g*. antibiotic clusters) are generally considered as ‘cryptic’ under most growth conditions^10^,^39^. We observed that they are poorly expressed in exponential phase. However their expression is characterized by up regulation after 48 h in the growth medium, compared to the rest of the chromosome including other genes putatively acquired by horizontal gene transfer (*e.g*. GIs). This result highlights that SMBGCs in *S. ambofaciens* have evolved regulatory mechanisms, such as the induction of cluster-situated transcriptional factor genes (**Supplementary Fig.2**) that efficiently and specifically regulate gene expression during metabolic differentiation, as reported in other *Streptomyces* species^4–7^.

Moreover, we show that changes in gene expression dynamics are correlated with metabolic differentiation. Indeed, the terminal regions become transcriptionally active after 48 h of growth (see also references^10^,^36^) (**Fig.2**). Interestingly, the ratio, antisense-over sense-transcription is high within the terminal compartments but decreases over the growth period, especially in the SMBGCs (**Extended Data Fig.2.F**). This suggests that this antisense-transcription may reflect regulatory processes that remain to be explored. This is particularly interesting since some NAPs suppress antisense-transcription in *S. venezuelae^40^*.

In addition, 3C-seq analysis revealed that in exponential phase the central compartment forms a multiple domain structure, delineated by boundaries, whereas the terminal regions form two large compartments in which contacts are more variable at larger distances. As previously shown for bacteria with circular genomes^20–23,26^, long and highly expressed genes (LHEGs encoding rRNA, ribosomal proteins, or respiratory chain components) are found at the boundaries in the *Streptomyces* chromosome (**Supplementary Table 5, Fig.4**). The positions of these boundaries are both conserved (under certain circumstances) and dynamic, since they change over the growth phase (^22^, this work). Notably, most of the boundaries observed at early growth times were correlated with the presence of persistent genes (**Fig.4C**). Within the central compartment, domains are a direct consequence of boundary formation. Additionally, no trivial role in gene expression or in chromosome conformation have been assigned to chromosome interacting domains (CIDs) in bacteria^41^. Here we propose that boundaries are likely to act as transcriptional hubs for clustered and persistent genes^42^, associated with very high levels of transcription in exponential phase. Moreover, in exponential phase, the first and last rDNA operons form the sharpest boundaries, recapitulating the borders that were arbitrarily used to define the central region beyond which genome synteny falls (**Figure 1**). We thus propose that they also constitute an evolutionary barrier that limits the occurrence of single recombination events within the central region. Interestingly, some boundaries in *Streptomyces* do not display high levels of gene expression (**Supplementary Table 5**). These are generally located within the terminal arms, suggesting the existence of other mechanisms, yet to be discovered that impose chromosome constraints.

Furthermore, we show for the first time that the central and terminal compartments present different organizational features, an observation independently demonstrated for the *S. venezuelae* chromosome (see accompanying paper, Szafran *et al*.). We favor the idea that the different organization of the terminal compartments could be a consequence of multiple factors: i) the lack of constraints imposed by active and sense-transcription; ii) the high level of anti-sense transcription; iii) the enrichment of horizontally acquired NAPs and other transcriptional regulators that, by increasing the dynamics of DNA intramolecular interactions, ensure appropriate SMBGC repression in exponential phase. Interestingly, it was shown in *S. venezuelae* that during sporulation the HupS NAP is involved in promoting optimal contacts along the whole chromosome, but seems to be particularly important for the organization of the terminal regions (see accompanying paper, Szafran *et al*.).

During metabolic differentiation, the changes in transcriptional dynamics are accompanied by a huge remodeling of chromosome folding, switching from an ‘open’ to a ‘closed’ conformation, in which highly expressed SMBGC genes form new boundaries (**Fig.4.E & F**). This may illustrate a relocation of transcriptional hubs from clustered and persistent genes (in exponential phase) to SMBGCs at the onset of metabolic differentiation. Remarkably, the central region is no longer structured into multiple domains. The lack of boundaries in the central compartment after 48 h growth is correlated with a slight decrease in transcripts from genes that were present in boundaries in exponential phase. Indeed most of these genes remain expressed at high level (‘CAT-3’ or ‘CAT_4’) but below a threshold classically associated with boundaries (a string of genes with above 20 000 normalized reads per kb, **Supplementary Table 5**). Moreover, we cannot exclude the possibility that some of these transcripts are stable RNAs produced in exponential rather than stationary phase^43^. This loss of boundaries in the central compartment is reminiscent of the phenomena observed in eukaryotes during G1-to-mitosis transition^44^ and could reflect a more general phenomenon of genome compaction during the cell cycle.

At larger scales, there is an increase in the frequency of inter-arm DNA contacts. The most likely candidate to be involved in this reorganization is the Smc-ScpAB condensin complex, which is recruited to the origin region by ParB bound to *parS^45,46^*. From the *parS* sites, Smc-ScpAB promotes inter-arm contacts by translocating to the terminal region of the chromosome, likely by loop extrusion^20,21,24,27,47–52^ *Streptomyces* spp. encode both a ParABS system^53–55^ and a Smc-ScpAB complex^56^. Interestingly, during sporulation *Streptomyces* chromosomes need to be compacted and segregated to ensure correct spore formation^57 56^ Indeed, HiC studies during sporogenic development in *S. venezuelae* showed that inter-arm contacts are dependent on Smc-ScpAB (Szafran *et al*., see accompanying paper). The global remodeling of the *Streptomyces* chromosome going from an ‘open’ to a ‘closed’ conformation seems to occur similarly during metabolic (this work) and sporogenic development (Szafran *et al*., see accompanying paper). Such a global rearrangement has not been previously described in wild-type bacteria. Interestingly, we observed a slight tilt in the secondary diagonal in contact analysis when it approaches to the terminal compartments (**Fig.4.E, Supplementary Fig.3.C**). This is consistent with a slow-down of Smc-ScpAB activity when interacting with the transcription machinery^50,58^, in agreement with the fact that transcription in the terminal compartment is higher than in the central compartment during metabolic differentiation (**Fig.4.E**, **Supplementary Fig.3.C**).

Collectively, these results indicate a link between evolutionary processes, including genome-compartmentalization and the molecular mechanisms (*e.g*. transcription, 3D-folding) that shape the structure and function of genes and genomes in *Streptomyces*. We therefore hypothesize that the efficiency of the regulatory processes controlling conditional expression of SMBGCs may be an emerging property of spatial compartmentalization. We believe that this study will open new insights into setting the rules governing chromosome spatial organization, expression and stability, and the optimal design of *Streptomyces* genomes for SMBCG expression.

## Methods

### *Streptomyces* strains and growth conditions

*S. ambofaciens* ATCC 23877 was grown on solid soy flour-mannitol (SFM) medium^59^ at 28°C unless otherwise indicated. The strain was grown in the following media: MP5 medium (7 g/l yeast extract, 20.9 g/l MOPS, 5 g/l NaCl, 1 g/l NaNO_3_, 36 ml/l glycerol – pH 7.5) ^34^, or YEME medium 10.3 % sucrose (3 g/l yeast extract, 5 g/l bactotryptone, 3 g/l malt extract, 10 g/l glucose, 103 g/l sucrose; pH 7.0-7.2; adapted from^59^). Twenty million spores of *S. ambofaciens* ATCC 23877 were inoculated in 100 ml of media before growth at 30°C in a shaking agitator (220 rpm, INFORS HT multitron standard).

### Bioassays

For bioassays from liquid cultures, 50 μl of filtered supernatants were spotted on to agar medium using cylinder (diameter 0.5 cm) and allowed to dry until complete penetration into a plate containing 50 ml of DIFCO antibiotic medium 5 (DIFCO 227710). Thereafter 7 ml of SNA medium (2 g/l agar, 8 g/l nutrient broth MP Biomedicals Cat#1007917) containing *Micrococcus luteus* (final OD_600nm_ 0.04) were overlaid on the plate and incubated at 37°C. The growth inhibition area was measured 24 h later.

### Genomic island (GI) identification

We designed the Synteruptor program (http://bim.i2bc.paris-saclay.fr/synteruptor/)^60^ to compare the sequences of chromosomes of species close enough to have synteny blocks, and to identify the genomic islands existing in each respective chromosome. We define synteny breaks as genomic regions between two consecutive synteny blocks when comparing two genomes, with the two blocks having the same direction. The genome of *S. ambofaciens* ATCC 23877 was compared to the chromosome of 7 closely related strains: *S. ambofaciens* DSM40697, *S. coelicolor* A3(2), *S. lividans* 1326, *S. lividans* TK24, *S. mutabilis, Streptomyces* sp. FXJ7.023, *Streptomyces* spp. M1013. To define a region harbouring GIs, we used a threshold of 15 CDS as the minimal number of CDSs within the synteny break, in at least one of the strains used for the pairwise comparison. Of note, a GI can therefore correspond to 15 CDS in one strain but less CDS in the other one. However, when the corresponding position within the *S. ambofaciens* genome contained less than 2 CDS or only tRNAs, compared to at least 15 CDSs in the chromosome of the compared species, it was considered as an insertion point (but not a GI) within *S. ambofaciens*. When the GIs were identified during several pairwise comparisons of *S. ambofaciens* ATCC 23877 and/or overlapping, they were fused and considered thereafter as a single GI. The complete list of GIs identified in the *S. ambofaciens* genome is presented in **Supplementary Fig. 1** and **Supplementary Table 2**.

### SMBGCs identification

We used antiSMASH5.1.0^33^ to identify putative SMBGCs. Thirty SMBGCs have thus been identified in the *S. ambofaciens* genome, the one encoding kinamycin biosynthesis being duplicated owing to its location within the TIRs. The definition of cluster boundaries has been manually refined on the basis of literature data for characterized SMBGCs (**Supplementary Table 4**). Of note, SMBGC genes are ≈ 1.5 times larger than the average, and only 13 of them (from 4 SMBGCs, namely ‘CL4_Indole’, ‘CL10_Furan’, ‘CL11_NRPS’, ‘CL20_Hopenoid’) belong to the core genome.

### NAPSFs, identification

In this study, we considered NAPs and chromosome structural factors *sensu lato* by including orthologues of classical NAPs and structural factors (HU, sIHF, Lsr2, Lrp/AsnC, SMC, Dps, CbpA, DnaA or IciA family proteins)^61^ and/or proteins associated with *S. coelicolor* chromatin^62^. This list is available in **Supplementary Table 5.**

### Definition of indexes for genome conservation analyses

We selected 125 *Streptomyces* genomes from the NCBI database representative of the *Streptomyces* genus by keeping only complete genomes of distinct species. When genomes share an average nucleotide identity (ANIb) value greater than or equal to 96 %, they are considered as members of the same species^31^ (**Supplementary Table 1**). We made one exception by keeping two strains of *S. ambofaciens* (ATCC 23877 and DSM 40697). Orthologous genes were identified by BLASTp best hits^63^ with at least 40 % of identity, 70 % coverage and an E-value lower than 1e^−10 31^. The gene persistence index was calculated as *N*_orth_/*N*, where *N*_orth_ is the number of genomes carrying a given orthologue and *N* the number of genomes searched^28^. Pairwise comparisons are achieved using a sliding window and comparing a reference strain (species A) to another (species B). Here is considered a window containing 8 genes noted from 1 to 8 in the reference species. To calculate the GOC index (Gene Order Conservation), which is the number of contiguous syntenic orthologs between the two chromosomes over the number of orthologs between the two chromosomes, calculated in a sliding window, the pair of contiguous genes present in the window in the reference species are searched and counted as contiguous pairs of orthologues in the whole genome of species B related to the number of orthologs is defined by the window.

### Transcriptome analysis

For RNA-seq analysis performed in liquid cultures, ≈ 2.10^7^ spores of *S. ambofaciens* ATCC 23877 were inoculated in 100 ml of liquid medium. Thereafter, 25 ml (for samples harvested after 24 h growth) or 10 ml of cultures (for other time points) were added to an equal volume of cold ethanol.

Cells were then harvested by centrifugation for 15 min at 4,000 *g* at 4°C, and stored at −20°C. Pellets were washed with 1 ml of DEPC water, centrifuged for 5 min at 16,000 *g* at 4°C and homogenized with glass beads (≤ 106 μm; G4649, SIGMA) in 350 μl of lysis buffer (RNeasy Plus Mini Kit, QIAGEN) supplemented with 10 μl/ml β-mercaptoethanol. Samples were processed 3 times for 45 sec each in FastPrep-24™ 5G instrument (MP Biomedicals) at setting 6 with 1 min cooling between the stages. After centrifugation for 10 min at 16,000 *g* at 4°C, total RNAs were isolated from the supernatants using an RNeasy Plus Mini Kit (QIAGEN) and gDNA Eliminator columns, following the manufacturer’s recommendations. To remove genomic DNA, RNA samples were incubated for 30 min at 37°C with 20 U of RNase-free DNase I (Roche) in a final reaction volume of 30 μl. RNAs were then cleaned up using the RNeasy Mini Kit (QIAGEN), following the manufacturer’s recommendations. The absence of DNA in the preparations was checked by PCR on an aliquot. RNA samples were quantified using Qubit™ RNA HS Assay kit (ThermoFischer Scientific), following manufacturer’s recommendations.

Total RNA quality was assessed in an Agilent Bioanalyzer 2100, using RNA 6000 pico kit (Agilent Technologies). 500 ng of total RNA were treated with DNAse (Baseline Zero DNAse, Epicentre) prior to ribosomal RNA depletion using the RiboZero bacteria magnetic Kit from Illumina according to the manufacturer’s recommendations. After the Ribo-Zero step, the samples were checked in the Agilent Bioanalyzer for complete rRNA depletion. Directional RNA-seq libraries were constructed using the Illumina ScriptSeq kit V2 (discontinued) for samples corresponding to C5 conditions and Illumina Stranded library preparation kit for all other samples, according to the manufacturer’s recommendations.

Libraries were pooled in equimolar proportions and sequenced (Paired-end 2×43 bp) with an Illumina NextSeq500 instrument, using a NextSeq 500 High Output 75 cycles kit. Demultiplexing was done (bcl2fastq2 V2.2.18.12) and adapters (adapter_3p_R1: AGATCGGAAGAGCACACGTCTGAACT; adapter_3p_R2: AGATCGGAAGAGCGTCGTGTAGGGA) were trimmed with Cutadapt1.15, only reads longer than 10 bp were kept.

STAR software ^64^ was used for mapping RNA-seq to the reference genome (genome-build-accession NCBI_Assembly: GCF_001267885.1) containing only one terminal inverted repeat (TIR). This avoids any biases with multiple mapping within the duplicated extremities of the genome (since the two TIR sequences are indistinguishable). We used the *featureCounts* program^65^ to quantify reads (in sense- and antisense-orientation) generated from RNA-sequencing technology in terms of “Gene” feature characteristics of *S. ambofaciens* ATCC 23877 most recent annotation (GCF_001267885.1_ASM126788v1_genomic.gff – released on the 06/15/2020).

### Bioinformatic analysis of RNA-seq count data

SARTools (Statistical Analysis of RNA-Seq data Tools) DESeq2-based R pipeline^66^ was used for systematic quality controls and the detection of differentially expressed genes. PCA and sample clustering used homoscedastic data transformed with Variance Stabilizing Transformation (VST). For the differential analysis, the Benjamini and Hochberg’s method was used with a threshold of statistical significance adjusted to 0.05. The reference condition was C1 (medium “MP5_24h”). All parameters of SARTools that have default values were kept unchanged. Genes with null read counts in the 37 samples were not taken into account for the analysis with DESeq2. Genes corresponding to the second TIR (not used for the mapping) or the rRNAs (RiboZero treatment) were excluded from further analysis. **Supplementary Table 5** presents the normalized counts per gene in each growth condition. To ensure that antisense-transcription did not affect the normalization of sense-transcription data, the latter were normalized as follows: the percentage of transcription in antisense orientation of each gene was determined with the raw data, then the normalized antisense-transcription was evaluated by applying this percentage to the normalized sense-transcript counts. To determine the category of gene expression the normalized counts obtained by the SARTools DESeq2-based pipeline were again normalized with respect to gene size (number of DESeq2 normalized reads/gene size x 1000 bp), and thereafter expressed in RPK (reads per kb). This allowed us to compare directly the relative levels of gene expression of individual genes. Data were analyzed with R software^67^ and the Integrative Genomics Viewer (IGV) tool was used to simultaneously visualize RNAseq data and genomic annotations^68^.

### Multidimensional analyses of the data

We used the FactoShiny R package^69^ to perform clustering, principal component analyses and correspondence analyses. This overlay factor map presented in **Extended Data Fig.2.C** results from the correspondence analysis performed using a contingency table indicating the number of genes in each category of expression level depending on the genome features of interest. Max’, ‘Mean’ and ‘Min’ refer to the maximal, mean and minimal expression levels in all studied condition, from the lowest category (‘0’) to the highest (‘4’). These categories were defined by considering the distribution parameters of the normalized number of reads (**Extended Fig.2.A.**). These gene expression categories as well as the number of genes switched ON (‘Switch’, meaning that the expression level switches from CAT_0 in at least one condition to CAT_3 or more in another condition), or presenting a very low (‘AS < 0.05’) or high (‘AS > 0.5’) level of antisense index were used to build the map, and projected on the plan. Here are some basic explanations of how to interpret the results of correspondence analysis. On **Extended Data Fig.2.C**, the squares correspond to the raw labels (genome features). The triangles correspond to the column labels (*e.g*. category of level of mean, maximal, minimal sense-transcription, antisense index < 0.05, antisense index > 0.5).

### Chromosome conformation capture (3C)

Spores of *S. ambofaciens* ATCC 23877 (≈ 4.10^7^) were inoculated in 200 ml of MP5 or YEME liquid medium. In these media, bacterial growth was monitored by opacimetry. At the indicated time point, cells were harvested from 100 ml samples of culture, adjusted to an OD_600nm_ of about 0.15 (with the appropriate fresh medium) and fixed by adding formaldehyde solution (3 % final concentration). Cells were then incubated under moderate agitation for 30 min at RT and 30 min more at 4°C. Glycine (250 mM final concentration) was added, and the bacteria incubated 30 min at 4°C with moderate agitation. Cells were then harvested by centrifugation for 10 min at 4,000 *g* at 4°C. The cells were gently suspended in 50 ml of PBS 1X and then again harvested by centrifugation for 10 min at 4,000 *g* at 4°C. This washing step was repeated once before suspending the cells in 1 ml of PBS before final harvesting by 10 min at 4,000 *g* at 4°C. The dry pellets were stored at −80°C until use.

Frozen pellets of exponentially grown cells were thawed, suspended in 600 μl Tris 10 mM EDTA 0.5 mM (TE) (pH 8) with 4 μl of lysozyme (35 U/μl; Tebu Bio) and incubated at RT for 20 min. For the samples collected after 48 h, the pellets were thawed, suspended in TE (pH 8) with 4 μl of lysozyme (35 U/μl; Tebu Bio) for 45 min and then homogenized with a Bioruptor sonication device (3 cycles of 30 seconds, with a pause of 30 second pause for each). Then, for both 24 h and 48 h samples, SDS was added to the mix (final concentration 0.5%) of cells and incubated for 10 minutes at RT. 500 μl of lysed cells were transferred to a tube containing 4.5 ml of digestion mix (1X NEB 3 buffer, 1% triton X-100) and 100 μl of the lysed cells were transferred to a tube containing 0.9 ml of digestion mix. 800 units of SalI were added to the 5 ml digestion mix. Both tubes were then incubated for 2 h at 37°C and 250 units of SalI were added to the 5 ml tube and further incubated, 2 h at 37°C. To stop the digestion reaction, 4 ml of the digestion mix were immediately centrifuged for 20 min at 20,000 *g*, and pellets suspended in 4 ml of sterile water. The digested DNA (4 ml in total) was split into 4 aliquots and diluted in 8 ml ligation buffer (1X ligation buffer NEB 3 (without ATP), 1 mM ATP, 0.1 mg/ml BSA, 125 Units of T4 DNA ligase 5 U/μl). Ligation was performed at 16°C for 4 h, followed by incubation overnight at 65°C with 100 μl of proteinase K (20 mg/ml) and 100μl EDTA 500 mM. DNA was then precipitated with an equal volume of 3 M Na-Acetate (pH 5.2) and two volumes of iso-propanol. After one hour at −80°C, DNA was pelleted and suspended in 500 μl 1X TE buffer. The remaining 1ml digestion mix with or without SalI were directly incubated with 100 μl of proteinase K (20 mg/ml) overnight at 65°C. Finally, all the tubes were transferred into 2 ml centrifuge tubes (8 tubes), extracted once with 400 μl phenol-chloroform pH 8.0, precipitated, washed with 1 ml cold ethanol 70% and diluted in 30 μl 1X TE buffer in the presence of RNAse A (1 μg/ml). Tubes containing the ligated DNA (3C libraries) were pooled. The efficiency of the 3C preparation was assayed by running aliquots of the 3C-librairies, the digested DNA or the non-digested DNA on 1% agarose gel. Finally the 3C libraries were quantified on the gel using QuantityOne software (BioRad).

### Processing of libraries for Illumina sequencing

Approximately 5 μg of a 3C library was suspended in water (final volume 130 μL) and sheared using a Covaris S220 instrument (Duty cycle 5, Intensity 5, cycles/burst 200, time 60 sec for 4 cycles). The DNA was purified using a Qiaquick^®^ PCR purification kit, DNA ends were prepared for adapter ligation following standard protocols^70^. Custom-made adapters^21^ were ligated overnight at 4°C. Ligase was inactivated by incubating the tubes at 65°C for 20 min. To purify DNA fragments ranging in size from 400 to 900 pb, a PippinPrep apparatus (SAGE Science) was used. For each library, one PCR reaction of 12 cycles was performed using 2 to 3 μl of 3C library, 0.2 μM Illumina primers PE1.0 and PE2.0 and 1 unit of Taq Phusion [Finnzymes]. The PCR product was purified on Qiagen MinElute columns and dimers of primers were removed from the 3C library by using AMPure XP beads following the manufacturer’s protocol (Beckman Coulter). Finally, libraries were subjected to 75 bp paired-end sequencing in an Illumina sequencer (NextSeq500).

### Processing of sequencing data

PCR duplicates from each 3C library sequence dataset were discarded using the 6 Ns of custom-made adapters^21^. Reads were aligned independently using Bowtie 2 in very sensitive mode^71^. Only reads with mapping quality > 30 were kept to establish contact maps.

### Generation of contact maps

Contact maps were built as described previously^72^. Briefly, each read was assigned to a restriction fragment. Non-informative events such as self-circularized restriction fragments, or uncut co-linear restriction fragments were discarded^73^. The chromosome of *S. ambofaciens* devoid of the TIRs was divided into 10 kbp bins and the frequencies of contacts between genomic loci for each bin were assigned. Contact frequencies were visualized as heatmaps. Raw contact maps were normalized with the sequential component normalization procedure (SCN^73^). To facilitate visualization, contact matrices are visualized as log matrices. First we applied to the SCN matrices the log10 and then a Gaussian filter (H=1) to smooth the image. The contact score for a pair of bins that due to mapping was identified as an outlier (z-score>contact score) was replaced with the median value of the contact matrix.

### Frontier Index determination

To analyze the domain organization of *S. ambofaciens*, we used the frontier index (FI) method that quantifies the involvement of each bin in the frontier of any domain, i.e. at any scale – see technical details in^37^. Briefly, this method consists, first, in computing derivatives of the contact maps. To this end, we used normalized contact maps where the average contact frequency, *P*(*s*), was substracted to reduce noise coming from the natural overall decrease of contact frequencies as the genomic distance (*s*) between loci increase. Here, we also added pseudocounts (equal to *m*/20 where *m* is the maximal value of the contact frequencies) such that all values of the contact maps are strictly positive and we considered the logarithm of the resulting maps in order to mitigate the strongest variations close to the diagonal. We then considered separately the derivatives of these maps along the vertical and horizontal axes, whose large positive values (in the upper part of the maps) are respectively associated with upstream and downstream frontiers of domains, respectively. After setting negative values to zero (to reduce noise even more), for each bin the corresponding upstream and downstream FIs were defined as the sum over all other bins at a distance below 600 kb of the resulting signals. We then computed the profile of upstream and downstream FIs. We considered peaks that are located above the median of the peak values plus two times the standard deviation (estimated by the 1.48 times the median absolute deviation to mitigate the impact of outliers). These statistics were computed separately for the left terminal arm, the central region and the right terminal arm as these compartments have distinct statistical 3C-seq features. Altogether, this led us to a list of significant upstream (orange) and downstream (green) peaks, respectively.

A boundary was then allocated to any bin (as well as to pairs of consecutive bins to cope with uncertainty of peak positions) for which both the upstream and downstream peaks were significant. To analyze the genetic environment associated with a boundary, we considered the environment around the boundary identified by the frontier index (± 2 bins). Note that due to intrinsically noisy nature of 3C-seq data, we detected bins with only an upstream or a downstream significant peak (**Supplementary Fig.3.D**). These bins with an ‘orphan’ peak were not associated with a boundary, and hence, were removed from the analysis.

### The dispersion index

The dispersion index of a signal reflects the range of variations relative to the mean value of the signal. Here, we compute it for the frequency of contacts. More precisely, it is defined as: *I*(*s*) = *var*(*s*)/*P*(*s*), where *P*(*s*) and *var*(*s*) are, respectively, the mean value and variance of the frequency of contacts between bins separated by the genomic distance. These quantities are then computed separately in each compartment of the genome. The larger the dispersion index is, the more variable are the contact frequencies within a compartment.

### Statistical procedure

Data were analyzed with R software^67^. For the RNA-seq analyses, three independent experiments (performed on different days) were carried out for each studied condition, except for C3, C9 and C10 that were performed in duplicate and C5 in quadruple. The statistical significance of RNA-seq analysis was assessed using the SARTools DESeq2-based pipeline^66^. For the 3C-seq analyses, two independent experiments (performed on different days) were carried out for each condition. To quantify gene regionalization, we defined the left and right terminal arms using as limits the first (genome position: 1,469,670-1,474,823 bp) and last (genome position: 5,802,471-5,807,629 bp) rDNA operons, respectively. The statistical significance was assessed by means of Fisher’s exact test for count data, which is appropriated to the analysis of contingency tables.

## Data availability

The datasets generated during this study have been deposited in the NCBI Gene Expression Omnibus (GEO, https://www.ncbi.nlm.nih.gov/geo/) under the accession number GSE162865.

## List of abbreviations

3C: Chromosome Conformation Capture
“Actino. Sign.”: Actinobacterial Signature
BGC: Biosynthetic Gene Cluster
CDS: coding sequence
CID: Chromosome Interacting Domain
GI: Genomic island
IGV: Integrative Genomics Viewer
LHEG: long and highly expressed genes
NAP: Nucleoid Associated Protein
NAPSFs: Nucleoid Associated Proteins and structural factors
PCA: Principal Component Analysis
rDNA: ribosomal RNA encoding DNA
RPK: Read per kb
SARTools: Statistical Analysis of RNA-Seq data Tools
SMBGC: Specialized Metabolite Biosynthetic Gene Cluster
TIR: Terminal inverted repeat
VST: Variance Stabilizing Transformation

## Acknowledgments

We acknowledge the High-throughput sequencing facility of I2BC for its sequencing and bioinformatics expertise. We thank Barry Holland and Christophe Possoz for careful reading of the manuscript and the members of F. B. and S. L. for fruitful discussions and advice.

---

**Extended Data Figure 1:**
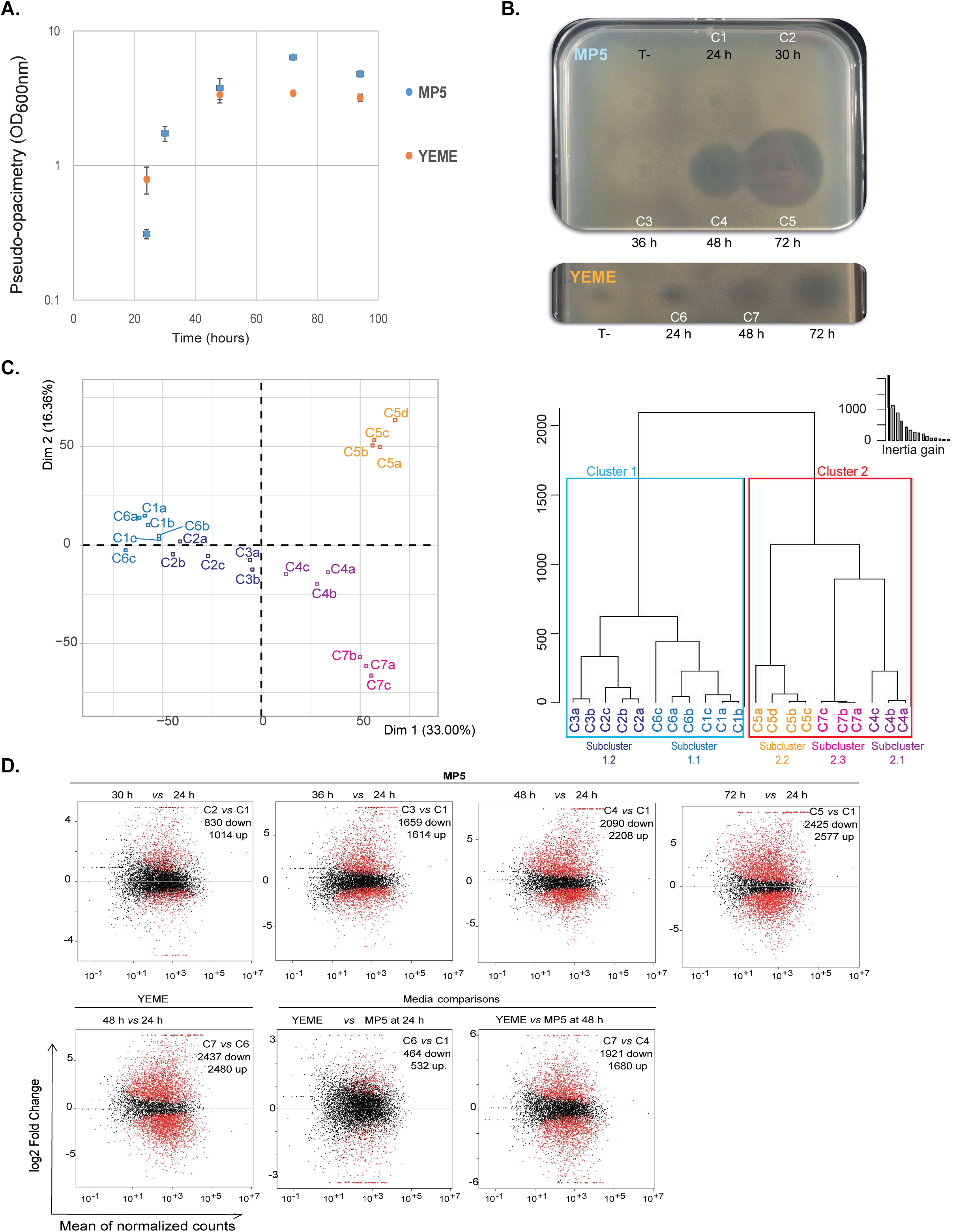
The metabolic differentiation of *Streptomyces ambofaciens* analyzed at the transcriptional level. A. **Growth curves of *Streptomyces ambofaciens* in MP5 and YEME liquid media.** In these media, *S. ambofaciens* grows under a rather dispersed way, so that growth can be monitored by optical density (OD_600nm_). We although refer to “pseudo-opacymetry” because of the presence of some clumps of bacteria. Values are means ± standard errors (error bars) from 3 independent experiments. B. **Bioassays performed in conditions used in this study.** Bioassays performed against *Micrococcus luteus* with the supernatants of *S. ambofaciens* ATCC 23877 grown in MP5 or YEME liquid media and harvested at different time points. Non-inoculated media were used as negative controls (“T-“). ‘C1’ to ‘C7’ refer to the name of the studied conditions (Table 1). The width of the plates is 12 cm. The presence of a halo indicates the presence of one or more antibacterial activities. The expression of the SMBGCs encoding known ATB is presented in **Supplementary Fig.2**. C. **Multidimensional analyses of the studied conditions. Left panel - Axes 1 and 2 of the principal component analysis (PCA)** of the whole data set (21 samples), with percentages of variance associated with each axis. The numbers indicate the condition (as indicated in Table 1) with a distinct letter (a, b, c, d) for each replica. **Right panel-Ascending hierarchical classification** (Euclidean distance, Ward criterion) and its inertia gain. The conditions form 2 clusters (1 and 2) and 5 subclusters which are boxed. The cluster 1 corresponds to the earliest growth time points (24 h, 30 h and 36 h) in which no ATB production was detected by bioassay. The cluster 2 corresponds to conditions associated to metabolic differentiation and the latest growth time points. Its sub-clusters separate owing the growth time and medium: 48 h in MP5 (subcluster 2.1), 72 h in MP5 (subcluster 2.2), 48 h in YEME (subcluster 2.3). D. **MA-plot of the data for the comparison of transcriptomes over growth and over media.** The MA-plot represents the log ratio of differential expression as a function of the mean intensity for each pairwise comparison. The differentially expressed features are highlighted in red, and the number of up- and down-regulated genes indicated on each graph. Triangles correspond to features having a too low/high log2(fold change) to be displayed on the plot.

**Extended Data Figure 2:**
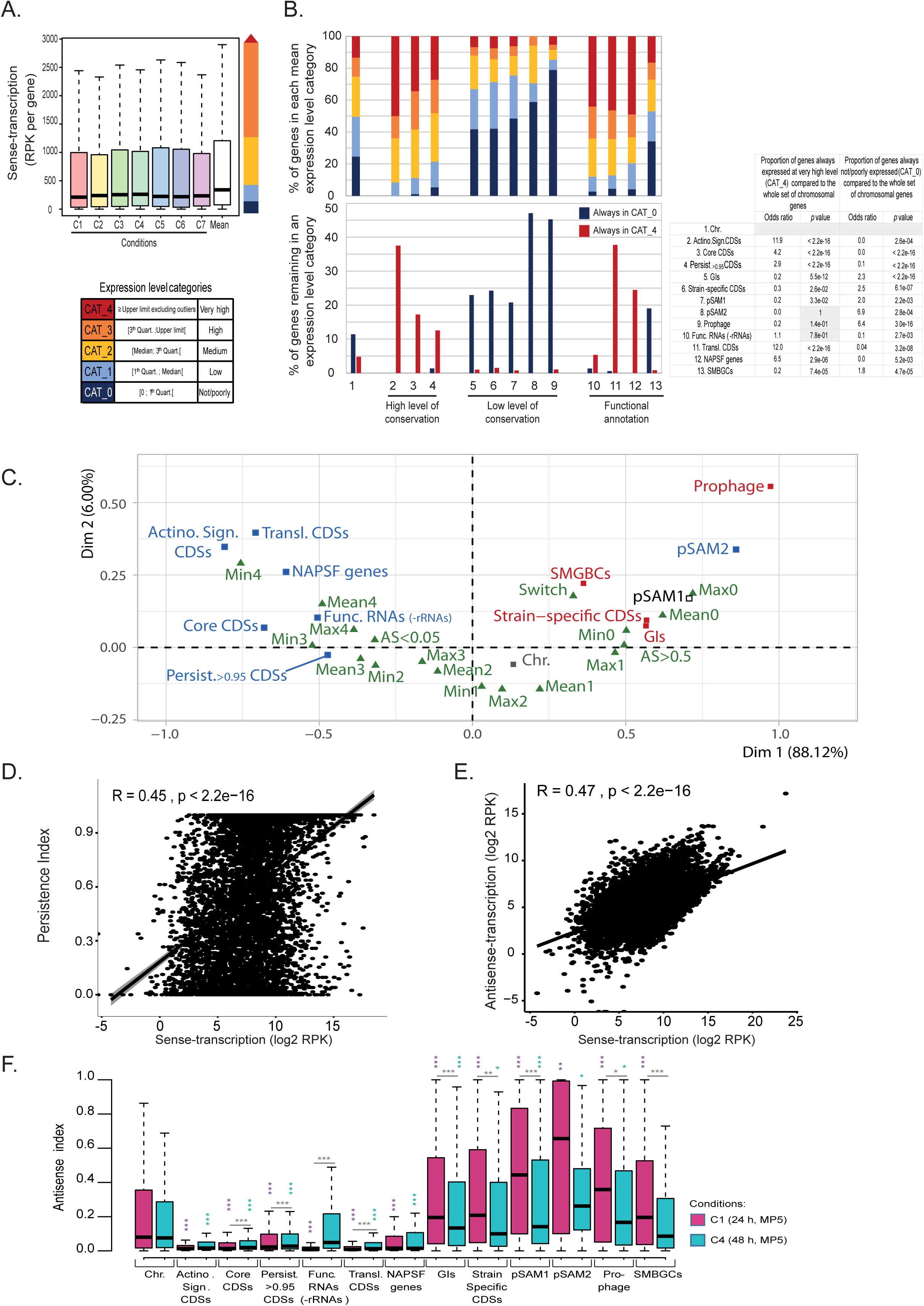
The dynamic of *Streptomyces ambofaciens* transcriptome. **A. Distribution of the normalized read counts per gene and definition of gene expression level categories.** The boxplot represents the distributions of the read counts per gene (normalized by DESeq and on gene size, RPK) individually for each tested condition and as the mean expression of each gene in all tested conditions. For clarity outliers were excluded from this graphical representation (but were taken into account for the numerical exploitation of the data). The genes were classified owing their expression level by considering the distribution parameters as indicated. These parameters were calculated both for each condition individually, and for the mean expression in all tested condition. **B. Pattern of gene expression depending on genome features of interest.** The genes were classified owing their mean (upper panel) or their minimal of maximal (lower panel) expression level in all studied conditions, as described in panel A. The table presents the statistical analysis of the data (Fisher’s exact test for count data). C. **Multidimensional analysis (overlay factor map) of genomic and transcriptomic data.** The squares and triangles correspond to genomic and transcriptomic features of interest, respectively (see abbreviations below). The closer the squares or triangles are to the origin of the graph, the closer they are likely to be to the mean data. Proximity between triangles or between squares indicates similarity. A small angle (*e.g*. less than 30°) connecting a triangle and a square to the origin indicate that they are probably associated. If they are located at the opposite, they are probably negatively associated. The proximity between the variables ‘SMBGCs’ and ‘Switch’ is due to the fact that a high proportion (19.3 %) of the SMBGC genes switched on at the transcriptional level in at least one studied condition (compared to 4.4 % at the level of the whole genome, odds ratios 5.2, *p*-value < 2.2.10^−16^; 7.6 % of genes within GIs, odds ratios 2.9, *p*-value 2.2.10^−9^, Fisher’s exact test for count data). <u>Genomic feature abbreviations</u>: ‘Actino. Sign. CDSs’ (coding sequences of the actinobacterial signature); ‘Chr.’ (whole chromosome); ‘Func. RNA (-rRNA)’ (Functional RNA excluded rRNA); GIs (genes belonging to genomic islands); ‘Persist.0.95 CDSs’ (coding sequences of *S. ambofaciens* ATCC 23877 presenting a gene persistence superior to 95 % in 124 other *Streptomyces* genomes); ‘Transl. CDSs’ (genes encoding functions involved in translation process and/or RNA stability); NAPSFs (nucleoid associated proteins and structural factors); SMBGCs (specialized metabolite BGCs). <u>Expression feature abbreviations</u>: ‘AS < 0.05’ and ‘AS > 0.5’ correspond to very low (below 0.05) or very high (above 0.5) antisense-transcription indices. ‘Max’, ‘Mean’ and ‘Min’ refer to the maximal, mean and minimal expression levels in all studied condition, from the lowest category (‘0’) to the highest (‘4’). ‘Switch’ correspond to the expression level switch from CAT_0 in at least one condition to CAT_3 or more in another condition. D. **Correlation between gene persistence and the transcription in sense orientation.** The correlation between gene persistence and the mean number of counts (normalized by DESeq2 and on gene size, RPK) in sense-orientation for each gene (over all studied conditions) was analyzed by a Spearman’s rank correlation test (not sensitive to the log2 conversion of the data). E. **Correlation between sense- and antisense-transcriptions.** The correlation between the mean number of counts (normalized by DESeq2 and on gene size, RPK) in sense- and antisense-orientation for each gene (over all studied conditions) was analyzed by a Spearman’s rank correlation test (not sensitive to the log2 conversion of the data). F. **Boxplot presenting the antisense index over growth in MP5 medium depending on feature of interest.** The distribution of the antisense index defined as the number of counts in antisense over total counts (in sense- and antisense-orientation) is presented depending on the genome features of interest (see panel C for abbreviations). For clarity outliers were excluded from this graphical representation (but were taken into account for the numerical exploitation of the data). The significant differences by Wilcoxon rank sum test with continuity correction are indicated by asterisks (*, *p*< 0.05; **, *p*< 0.01; ***, *p*< 0.001), in grey when comparing results after 24 h *versus* 48 h of growth for a given genome feature of interest, in violet and green when comparing a genome feature of interest to the whole set of chromosomal genes after 24 h *versus* 48 h of growth, respectively.

**Extended Data Figure 3:**
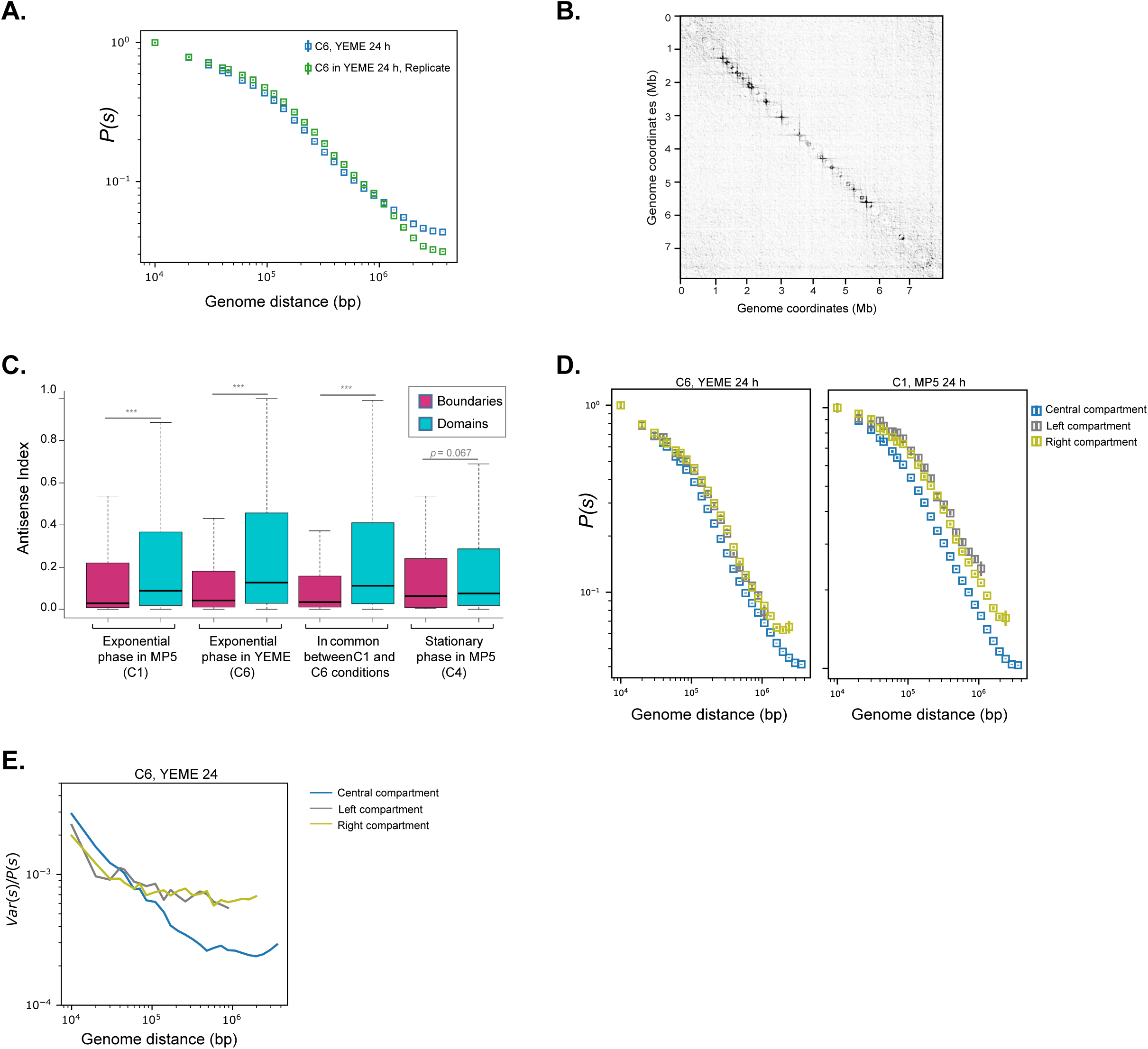
Data exploration linking the chromosome architecture, transcription and gene persistence. A. **Contact probability, *P*(*s*), plotted as a function of genomic distance in exponential phase in YEME medium (C6 condition)**. The plot represent the intra-chromosomal contact probability, *P*(*s*), for pairs of loci separated by a genomic distance (in base pairs) on the chromosome, showing that the probability of contact extends up to approximately 1 Mb. The results of two replicates (#1 and #2) are presented. B. **Derivative of the contact matrix for exponentially growing WT cells**. The map shows for each genome locus the tendency of contact frequencies to go downwards or upwards. Boundaries are visualized as the black vertices of the squares along the diagonal. Regions with no upstream/downstream bias show no squares (*e.g*. terminal regions). C. **Boxplot presenting the antisense index over growth depending of genes depending on their architectural location.** The distribution of the antisense index defined as the number of counts in antisense over total counts (in sense- and antisense-orientation) is presented depending on the location of the genes (within boundaries or domains). For clarity, outliers were excluded from this graphical representation (but were taken into account for the numerical exploitation of the data). The significant differences by Wilcoxon rank sum test with continuity correction (boundaries *versus* domains) are indicated by asterisks (***, *p*< 0.001). D. **Contact probability, *P*(*s*), of the left, central and right compartments plotted as a function of genomic distance**. The plot represents the intra-chromosomal contact probability, *P*(*s*), for pairs of loci separated by a genomic distance (in base pairs) on the three compartments. E. **Dispersion index, *Var*(*s*)/*P*(*s*), of the left, central and right compartments plotted as a function of genomic distance in YEME after 24 h of growth (C6 condition).** The index of dispersion (Var(s)/P(s)) reflects the range of variations relative to the mean value as a function of the genome distance for the contact maps obtained for cells grown 24 h in YEME medium. Long-range DNA contacts within the terminal compartments (> 100 kbp) are more variable than within the central compartment.

**Supplementary Figure 1: Size distribution of *S. ambofaciens* ATCC 23877 GIs** Using a minimal threshold of 15 CDS within the synteny break in at least one of the compared genome, 50 GIs (composed of 2 to 158 CDSs) and 22 insertion points (≤ 1 CDS within the synteny break) were identified in *S. ambofaciens* ATCC 23877 genome.

**Supplementary Figure 2: Transcriptomes over growth of the four biosynthetic gene clusters encoding all known antibacterial activities of *Streptomyces ambofaciens* ATCC 23877**

The genes expressed at the highest level is boxed in red for the congocidine, spiramycin and kinamycin BGCs. They correspond to *cgc1* (SAM23877_RS39345, encoding a transcriptional regulator), *srmB* (SAM23877_RS26680, encoding the ABC-F type ribosomal protection protein SrmB) and *alpZ* (SAM23877_RS00890, encoding a transcriptional regulator) genes, respectively. The gene encoding SrmS (SAM23877_RS26595), the transcriptional regulator of the spiramycin BGC, is boxed in black. The dots are joined by a line for clarity (but the lines do neither report a real observation nor a model).

**Supplementary Figure 3: 3C-contact-maps for *S. ambofaciens* grown in the studied conditions**

**A. Replicate of cells after 24 h in YEME growth medium, C6 condition**

**B. Replicate of after 24 h in MP5 growth medium, C1 condition**

**C. Replicate of after 48 h in MP5 growth medium, C4 condition.** The violet and pink arrows indicate the location of boundaries formed at desferrioxamin and congocidine BGCs, respectively.

**D. Results of the frontier index analysis including peaks that do not form boundaries**. This panel presents the frontier index analysis of the 3C-maps presented in **Fig.3** and **Fig.4**, showing single peaks that where not considered as indicative of a boundary. At each locus, two indices are computed, reflecting the intensity of the loss of contact frequencies when going downwards (green peaks) or upwards (orange peaks), respectively, to the locus. A boundary is defined as any bin in which there is a change in the right bias of contacts towards the left bias (± 2 bin, green and orange peaks, respectively). The detection of bins with only an upstream or a downstream significant peak reflects the intrinsically noisy nature of 3C-seq data.

**Supplementary table 1: Collection of 125 *Streptomyces* genomes used in this study**

**Name**: Species name

**Genome Assembly ID**: Unique identifier used in the NCBI Genome Assembly database

**Chromosome length (bp)**: Chromosome length expressed in bp

**Plasmid length (bp)**: Cumulative length of plasmids [Number of plasmids] expressed in bp

**CDSs nb**: Number of CDSs in chromosome [CDSs in plasmids]

**TIR length (bp)**: Size of one copy of a TIR expressed in bp.

**Supplementary table 2: List of the GIs identified in *S. ambofaciens* ATCC 23877 genome**

**Supplementary table 3: Gene distribution in the central region *versus* the extremities (defined by the first and last rDNA operons) of *Streptomyces ambofaciens* ATCC 23877 chromosome depending on features of interest**

**Supplementary table 4: List of the SMBGCs identified in *S. ambofaciens* ATCC 23877 genome**

**Supplementary table 5: Overall results from the comparative genomics, RNA-seq and 3C-seq analyses**

**ID:** Gene identifier (NCBI annotation).

**SUPPORT:** genetic support, either chromosome (CHR) or pSAM1 plasmid.

**TIR:** ‘TIR’ if the gene is located within a TIR, ‘NA’ otherwise.

**START:** start position of the gene on the genetic support.

**END:** end position of the gene on the genetic support.

**STRAND:** orientation of the gene on the genetic support.

**REPLICATION_ORIENTATION:** orientation of the gene regarding the origin of replication, either on the “leading” or the “lagging” strand.

**GENE_SIZE:** size of the gene (in bp).

**Protein_ID:** Protein identifier (NCBI annotation).

**GO_ID:** Gene product identifier (Uniprot or RNA Central, suitable for GO analysis).

**Gene_Name**: Gene name (from NCBI and manual annotation from SMBGCs encoding known antibiotics).

**UNIPROT_Gene_Name:** Uniprot gene name, old locus tag.

**Product (NCBI annotation):** Product annotation from NCBI.

**Product (Uniprot annotation):** Product annotation from Uniprot.

**ACTINO_SIGN:** ‘ACTINO_SIGN’ if the gene encodes a protein of the actinobacterial signature, ‘NA’ otherwise.

**CORE:** ‘CORE’ if the gene belongs to the core CDSs identified by comparing 125 *Streptomyces* genomes (see Methods section for details, as well as Supplementary Table 1).

**PERSIST95:** ‘PERSIST95’ if the gene persistence is superior to 95 %, ‘NOT_PERSIST95’ otherwise.

**GI:** annotation as to whether or not the gene belongs to a GI (see details about GI names on Supplementary Table 2).

**Strain-specific_CDSs:** ‘UNIQ’ if the gene is only present in *S. ambofaciens* ATCC23877 (in a panel of 125 *Streptomyces* genomes described in Supplementary Table 1), ‘NOT_UNIQ’ otherwise.

**Func_RNA:** annotation as to whether or not the gene encodes a functional RNA (tRNA, tmRNA, SRP RNA, RNAse P RNA) excluding rRNA.

**Transl._CDS**: ‘translation’ if the CDS encodes a function related to translation and/or RNA stability, ‘NA’ otherwise

**NAPSF**: ‘NAPSF’ if the gene encodes a nucleoid-associated protein or a chromosome structural protein and/or a protein experimentally found associated to *Streptomyces* chromatin (see Methods for details), ‘NA’ otherwise.

**SMBGC**: annotation as to whether or not the gene belongs to a SMBGC (see details about SMBGC names on Supplementary Table 4).

**pSAM2**: ‘pSAM2’ if the gene belongs to pSAM2 integrative plasmid, ‘NA’ otherwise. **PROPHAGE**: ‘PROPHAGE’ if the gene belongs to the prophage integrated in *S. ambofaciens* ATCC23877 genome, ‘NA’ otherwise

**PERSIST_INDEX**: Persistence index (*N*orth/*N*, where *N*orth is the number of genomes carrying a given orthologue and *N* the number of genomes searched, 124 in our case). This persistence index was calculated only for CDSs (functional RNAs were not considered).

**<u>‘SENSE’ columns</u>:** Number of normalized counts (DESeq2 normalization and normalized on gene size) per kb for each gene in the sense orientation, as the mean of the number of counts in all tested conditions (‘SENSE_Mean’) or of the replicates in each condition considered individually (‘SENSE_C1’ to ‘SENSE_C7’)

**<u>‘ANTISENSE INDEX’ columns</u>:** Number of reads in the antisense orientation over the total (sense plus antisense) number of reads for each gene in all tested conditions (‘ANTISENSE_INDEX_Mean’) or in each condition considered individually (‘ANTISENSE_INDEX _C1’ to ‘ANTISENSE_INDEX _C7’)

**<u>‘DIFF’ columns</u>:** annotation as to whether or not the gene is statistically differentially expressed compared to C1 condition, with an adjusted *p* value ≤ 0.05 (‘DIFF0.05’), 0.01 (‘DIFF0.01’) or 0.001 (‘DIFF0.001’).

**<u>‘CAT’ columns</u>:** Gene expression level category in all tested conditions (‘CAT _Mean’), in each condition considered individually (‘CAT _C1’ to ‘CAT _C7’), maximal category in all tested conditions (‘CAT_MAX’) and minimal category in all tested conditions (‘CAT_MIN’).

**SWITCH**: annotation as to whether or not the gene expression level switches from CAT_0 to a maximal CAT_3 (‘SWITCH0->3’) or CAT_4 (‘SWITCH0->4’) category, when considering all studied conditions.

**‘BORDER’ columns:** annotation as to whether or not the gene belongs to a boundary observed on the map contact obtained in C1 (‘C1_BORDER’), C4 (‘C4_BORDER’) and/or C6 condition (‘C4_BORDER’). The boundaries are numbered according to their order in the genome. ‘rRNA’ indicates that the boundary also contains an rDNA operon.

