## Supplementary figures and images for "Dynamics of the compartmentalized *Streptomyces* chromosome during metabolic differentiation"

### Supplementary_Figure_1

Supplementary Figure 1

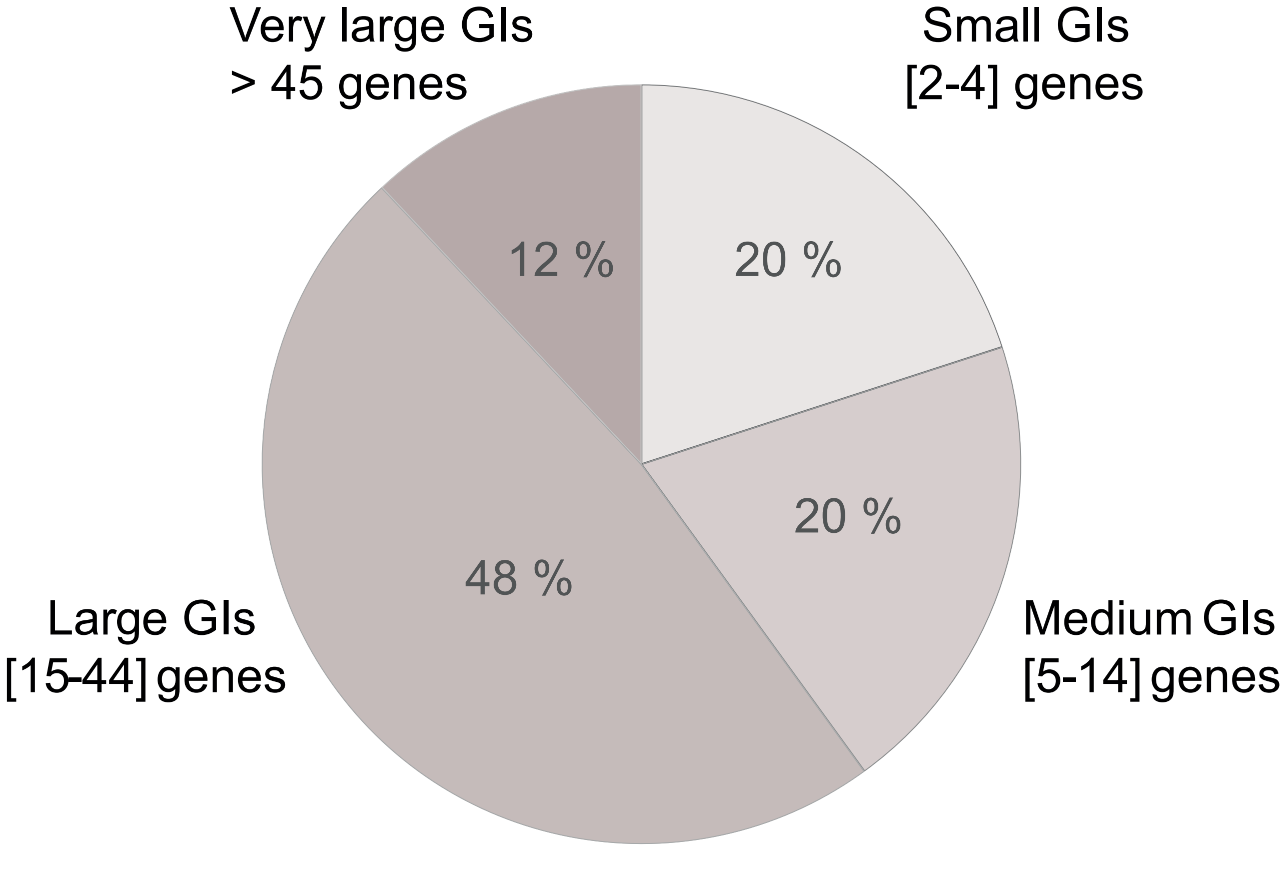

### Supplementary_Figure_2

#### Supplementary Figure 2

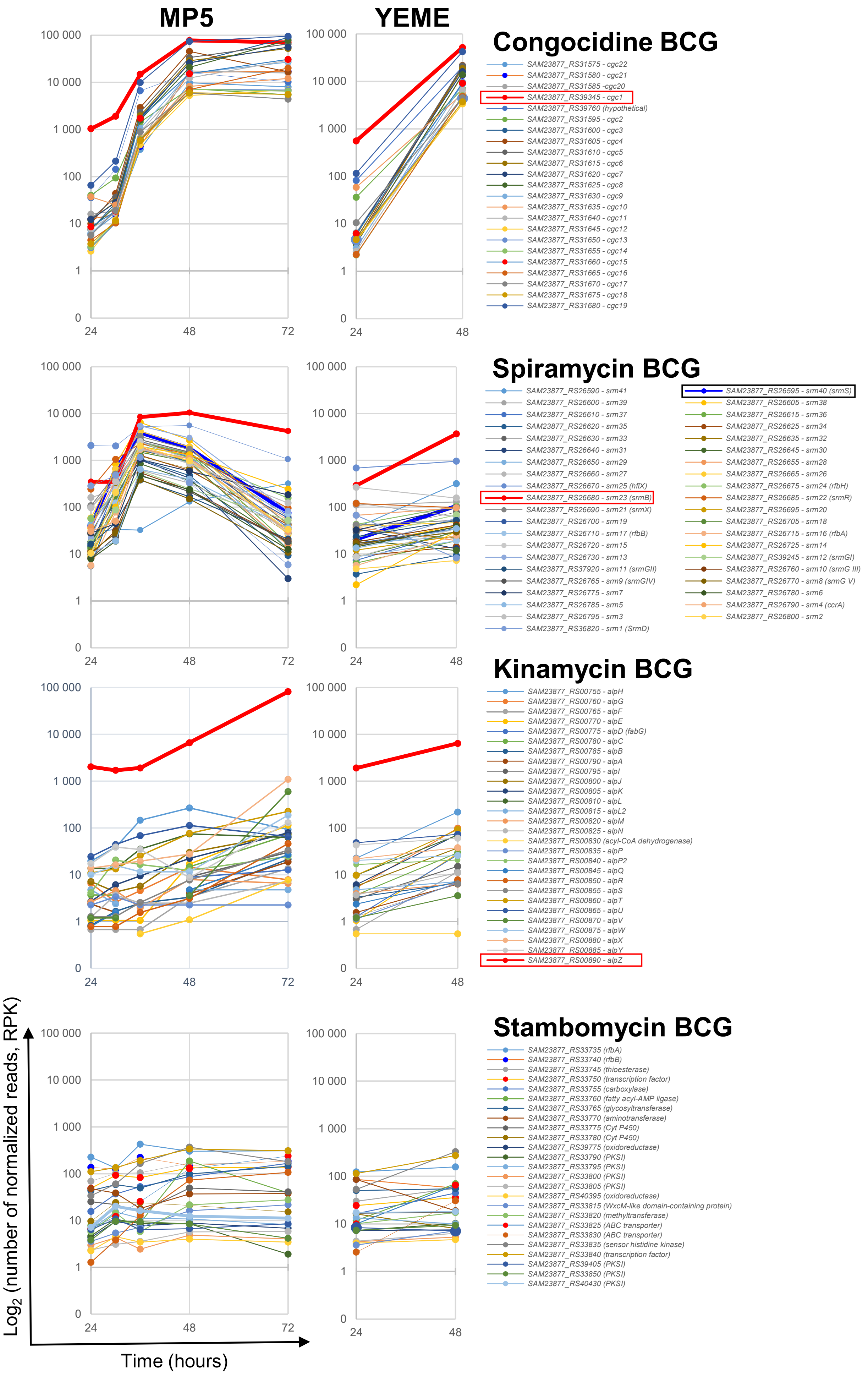

### Supplementary_Figure_3

##### Supplementary Figure 3

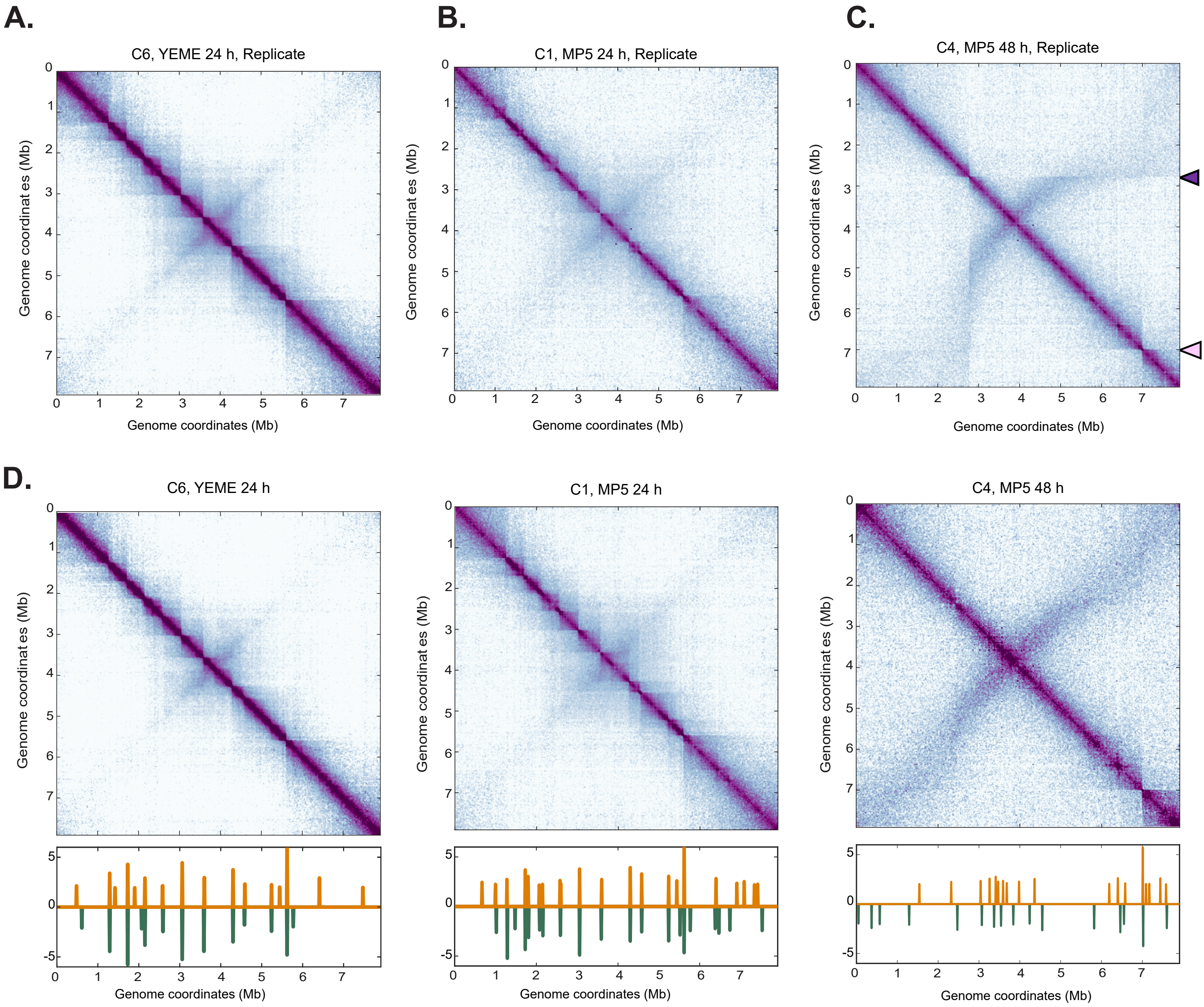
