## Supplementary_Table_1 for "Dynamics of the compartmentalized *Streptomyces* chromosome during metabolic differentiation"

| Name | Genome Assembly ID | Chromosome length (bp) | Plasmid length (bp) | CDSs nb | TIR length (bp) |
|---|---|---|---|---|---|
| *Streptomyces* sp. S1A1 7 | ASM711356v2 | 11,713,151 | 292,353 [1] | 11,068 [265] | ND |
| *S. bingchenggensis* BCW 1 | ASM9238v1 | 11,936,683 | - | 10,216 | ND |
| *Streptomyces* sp. 11 1 2 | ASM359554v1 | 11,603,877 | 51,329 [1] | 9,986 [61] | 29,227 |
| *Streptomyces* sp. GGCR 6 | ASM395571v1 | 10,393,987 | - | 9,540 | 96,011 |
| *Streptomyces* sp. CdTB01 | ASM148456v1 | 9,902,731 | 288,836 [1] | 9,444 [287] | 400,291 |
| *Streptomyces* sp. YIM 121038 | ASM608871v1 | 10,130,554 | 967,890 [1] | 9,440 [840] | 128,054 |
| *S. hygroscopicus jinggangensis* 5008 | ASM24535v1 | 10,145,833 | 237,851 [2] | 9,401 [243] | ND |
| *Streptomyces* sp. GY16 | ASM918486v1 | 10,116,136 | 59,304 [1] | 9,248 [66] | ND |
| *Streptomyces* sp. 769 | ASM81602v1 | 10,100,774 | 237,512 [1] | 9,180 [224] | 155,968 |
| *Streptomyces* sp. T44 | ASM979628v1 | 9,735,779 | - | 9,126 | ND |
| *S. scabiei* 87 22 | ASM9130v1 | 10,148,695 | - | 9,009 | 18,488 |
| *Streptomyces* sp. P3 | ASM303247v1 | 9,851,971 | - | 8,820 | ND |
| *S. noursei* ATCC 11455 | ASM170427v1 | 9,815,884 | - | 8,811 | 170,209 |
| *S. kanamyceticus* ATCC 12853 | ASM870449v1 | 10,133,897 | - | 8,806 | ND |
| *S. chartreusis* ATCC 14922 | ASM870471v1 | 9,912,098 | - | 8,704 | ND |
| *S. atratus* SCSIO ZH16 | ASM333086v1 | 9,641,288 | - | 8,700 | 288,246 |
| *S. lincolnensis* LC G | ASM334444v1 | 9,513,637 | - | 8,622 | ND |
| *S. griseorubiginosus* 3E 1 | ASM359523v1 | 9,512,378 | - | 8,551 | 113,251 |
| *S. lydicus* A02 | ASM95203v2 | 9,307,519 | - | 8,534 | 52,505 |
| *Streptomyces* sp. CC0208 | ASM344373v1 | 9,320,089 | - | 8,531 | ND |
| *S. coeruleorubidus* ATCC 13740 | ASM870513v1 | 9,335,698 | - | 8,518 | ND |
| *Streptomyces* sp. NEAU S7GS2 | ASM317327v1 | 9,641,634 | 45,805 [1] | 8,508 [52] | ND |
| *S. ectabilis* ATCC 27465 | ASM870479v1 | 9,807,160 | - | 8,329 | 112,273 |
| *S. coelicolor* A3 2 | ASM20383v1 | 8,667,507 | 387,340 [2] | 8,320 [400] | 21,653 |
| *S. formicae* KY5 | ASM255654v1 | 9,611,874 | - | 8,317 | 35,482 |
| *S. vietnamensis* GIM4 0001 | ASM83000v1 | 8,867,142 | 286,635 [1] | 8,287 [271] | 49,821 |
| *S. rimosus* ATCC 10970 | ASM870465v1 | 9,361,154 | - | 8,287 | 11,287 |
| *S. autolyticus* CGMCC0516 | ASM198397v1 | 10,029,028 | 155,632 [7] | 8,279 [36] | ND |
| *S. griseorubiginosus* BTU6 | ASM651693v1 | 9,226,027 | - | 8,274 | 224,191 |
| *S. albulus* CK 15 | ASM93518v3 | 9,336,218 | - | 8,241 | ND |
| *Streptomyces* sp. S10 2016 | ASM161179v1 | 9,083,372 | - | 8,241 | 134,209 |
| *S. dengpaensis* XZHG99 | ASM294683v1 | 8,541,354 | 168,817 [2] | 8,236 [210] | 21,003 |
| *Streptomyces* sp. ADI95 16 | ASM372149v1 | 8,184,000 | 902,421 [4] | 8,223 [861] | ND |
| *S. avermitilis* MA 4680 | ASM976v2 | 9,025,608 | 94,287 [1] | 8,132 [94] | ND |
| *S. lavendulae* CCM 3239 | ASM280384v1 | 8,691,711 | 241,081 [1] | 8,085 [231] | 237,734 |
| *S. chartreusis* NRRL 3882 | NRRL3882 | 8,983,317 | - | 8,083 | 39,831 |
| *S. ectabilis* NRRL 2792 | ASM736339v1 | 9,505,665 | - | 8,055 | ND |
| *Streptomyces* sp. M2 | ASM410450v1 | 8,718,751 | - | 8,025 | ND |
| *Streptomyces* sp. SYP A7193 | ASM949565v1 | 8,193,527 | 629,255 [2] | 7,984 [571] | ND |
| *S. lydicus* WYEC 108 | ASM399437v1 | 9,125,666 | - | 7,970 | 8,910 |
| *S. venezuelae* ATCC 14584 | ASM864231v1 | 8,942,078 | - | 7,935 | 165,025 |
| *S. lunaelactis* MM109 | ASM305455v1 | 8,396,100 | 174,091 [2] | 7,873 [149] | ND |
| *Streptomyces* sp. Mg1 | ASM41226v2 | 7,868,178 | 848,015 [3] | 7,864 [812] | ND |
| *Streptomyces* sp. QMT 28 | ASM949827v1 | 8,880,330 | - | 7,861 | ND |
| *S. griseoviridis* F1 27 | ASM399439v1 | 8,963,414 | - | 7,785 | 58,707 |
| *S. venezuelae* ATCC 15068 | ASM864237v1 | 8,558,202 | - | 7,743 | ND |
| *Streptomyces* sp. Go 475 | ASM333084v1 | 8,570,609 | - | 7,729 | 6,902 |
| *S. gilvosporeus* F607 | ASM208219v1 | 8,482,298 | - | 7,624 | ND |
| *S. olivoreticuli* ATCC 31159 | ASM339113v1 | 8,809,793 | - | 7,593 | 170,471 |
| *S. hundungensis* BH38 | ASM362781v1 | 8,393,044 | - | 7,541 | 65,122 |
| *S. ambofaciens* ATCC 23877 | ASM126788v1 | 8,303,940 | 89,658 [1] | 7,541 [128] | 202,695 |
| *S. alfalfae* ACCC40021 | ASM197502v1 | 8,625,867 | - | 7,525 | 149,241 |
| *Streptomyces* sp. SUK 48 | ASM965076v1 | 8,341,671 | - | 7,507 | 188,470 |
| *Streptomyces* sp. CCM MD2014 | ASM77204v1 | 8,274,043 | - | 7,501 | 14,382 |
| *S. pristinaespiralis* HCCB 10218 | ASM127807v1 | 8,532,592 | - | 7,498 | 419,828 |
| *S. platensis* ATCC 23948 | ASM870485v1 | 8,501,012 | - | 7,490 | 24,984 |
| *S. rochei* 7434AN4 | ASM806499v1 | 8,364,802 | - | 7,485 | 53,905 |
| *Streptomyces* sp. W1SF4 | ASM395003v1 | 7,272,878 | 795,265 [2] | 7,420 [656] | ND |
| *S. actuosus* ATCC 25421 | ASM320803v1 | 8,145,579 | - | 7,399 | 17,181 |
| *S. endophyte* N2 | ASM410448v1 | 8,428,700 | - | 7,398 | 53,557 |
| *Streptomyces* sp. Sge12 | ASM208045v1 | 7,983,613 | 127,085 [1] | 7,376 [119] | 162,199 |
| *S. pactum* ACT12 | ASM200522v1 | 8,550,793 | - | 7,375 | 239,610 |

| | | | | | |
|---|---|---|---|---|---|
| *Streptomyces* sp. WAC 01438 | ASM394552v1 | 8,138,328 | 62,839 [1] | 7,350 [72] | 49,769 |
| *S. lydicus* GS93 23 | ASM198444v1 | 8,243,179 | - | 7,320 | 36,373 |
| *S. griseus* NBRC 13350 | ASM1060v1 | 8,545,929 | - | 7,306 | 132,910 |
| *S. venezuelae* ATCC 14583 | ASM864235v1 | 8,018,484 | - | 7,301 | 157,265 |
| *Streptomyces* sp. SS52 | ASM479571v1 | 8,184,045 | - | 7,293 | 43,391 |
| *S. collinus* Tu 365 | ASM44487v1 | 8,272,925 | 104,361 [2] | 7,283 [101] | 631,364 |
| *Streptomyces* sp. WAC 01529 | ASM394554v1 | 8,270,461 | - | 7,269 | 58,922 |
| *Streptomyces* sp. KPB2 | ASM395005v1 | 8,082,236 | - | 7,256 | 40,616 |
| *S. ambofaciens* DSM 40697 | ASM163286v1 | 8,137,876 | - | 7,250 | 212,696 |
| *Streptomyces* sp. fd1 xmd | ASM200768v1 | 7,929,999 | - | 7,236 | ND |
| *S. clavuligerus* ATCC 27064 2 3 | ASM551946v1 | 6,748,591 | 1,795,495 [1] | 7,226 [1,534] | ND |
| *Streptomyces* sp. GSSD 12 | ASM334496v1 | 8,454,852 | - | 7,201 | ND |
| *S. katrae* S3 | ASM202842v1 | 7,504,851 | 551,599 [2] | 7,196 [321] | ND |
| *Streptomyces* sp. CFMR 7 CFMR 7 | ASM127809v1 | 8,207,742 | 99,537 [1] | 7,190 [95] | 11,739 |
| *S. cattleya* NRRL 8057 | ASM23730v1 | 6,283,062 | 1,809,491 [1] | 7,189 [1,651] | ND |
| *S. tsukubensis* AT3 | ASM929602v1 | 8,615,214 | 18,766 [1] | 7,183 [13] | 17,935 |
| *S. venezuelae* ATCC 14585 | ASM864233v1 | 8,048,154 | - | 7,176 | 166,199 |
| *S. leeuwenhoekii* sleC34 | sleC34 | 7,903,895 | 218,596 [2] | 7,120 [263] | 388,272 |
| *Streptomyces* sp. QHH 9511 | ASM978963v1 | 7,524,079 | 91,197 [1] | 7,109 [124] | ND |
| *S. niveus* SCSIO 3406 | ASM200917v1 | 7,990,492 | - | 7,105 | ND |
| *Streptomyces* sp. CB09001 | ASM336979v1 | 7,787,608 | - | 7,085 | ND |
| *S. globisporus* C 1027 | ASM26134v2 | 7,608,611 | 174,988 [2] | 7,079 [154] | ND |
| *S. fulvissimus* DSM 40593 | ASM38594v1 | 7,905,758 | - | 7,072 | ND |
| *S. albireticuli* MDJK11 | ASM219245v1 | 8,144,417 | - | 7,024 | 133,288 |
| *S. vinaceus* ATCC 27476 | ASM870493v1 | 7,673,509 | - | 7,020 | ND |
| *S. venezuelae* ATCC 10595 | ASM870525v1 | 7,871,480 | - | 7,001 | ND |
| *S. galilaeus* ATCC 14969 | ASM870457v1 | 7,756,194 | - | 7,001 | ND |
| *Streptomyces* sp. Tue6075 | ASM193163v1 | 7,931,832 | - | 6,994 | 12,088 |
| *S. alboniger* ATCC 12461 | ASM870439v1 | 7,962,786 | - | 6,993 | ND |
| *S. nigra* 452 | ASM307405v1 | 7,641,029 | - | 6,992 | 107,944 |
| *S. violaceoruber* S21 | ASM208217v1 | 7,916,045 | - | 6,979 | 4,122 |
| *S. parvulus* 2297 | ASM166004v1 | 7,149,446 | 617,085 [1] | 6,963 [443] | ND |
| *S. bacillaris* ATCC 15855 | ASM326867v1 | 7,888,441 | - | 6,953 | ND |
| *Streptomyces* sp. TN58 | ASM194184v1 | 7,585,034 | - | 6,936 | 193,939 |
| *S. venezuelae* ATCC 21018 | ASM864227v1 | 7,746,267 | - | 6,919 | ND |
| *S. nodosus* ATCC 14899 2 | ASM870499v1 | 7,772,587 | - | 6,917 | ND |
| *S. albus* ZD11 | ASM367534v1 | 8,317,371 | - | 6,904 | 614,456 |
| *S. subrutilus* ATCC 27467 | ASM870453v1 | 7,604,974 | - | 6,839 | 162,000 |
| *S. fungicidicus* TXX3120 | ASM38594v1 | 6,740,768 | 926,728 [1] | 6,839 [841] | ND |
| *S. tsukubensis* NRRL 18488 | ASM393271v1 | 7,963,742 | 55,806 [2] | 6,809 [63] | 210,577 |
| *S. cinereoruber* ATCC 19740 | ASM929938v1 | 7,516,652 | - | 6,807 | ND |
| *S. prasinus* ATCC 13879 | ASM870444v1 | 7,647,592 | - | 6,793 | ND |
| *Streptomyces* sp. SM18 | ASM291077v2 | 7,703,166 | - | 6,783 | 14,611 |
| *Streptomyces* sp. PAMC26508 | ASM36480v1 | 7,526,197 | 104,048 [1] | 6,779 [84] | 36,502 |
| *S. nitrosporeus* ATCC 12769 | ASM870455v1 | 7,581,562 | - | 6,778 | 26,990 |
| *Streptomyces* sp. SSL 25 | ASM785615v1 | 8,146,484 | - | 6,768 | 132,197 |
| *S. glaucescens* GLA O | ASM76121v1 | 7,453,200 | 170,574 [1] | 6,715 [140] | 14,128 |
| *S. asterosporus* DSM 41452 | ASM671613v1 | 7,766,581 | - | 6,710 | ND |
| *S. venezuelae* ATCC 21782 | ASM864229v1 | 7,525,322 | - | 6,699 | 182,221 |
| *S. cyaneogriseus* noncyanogenus NMWT 1 | ASM93144v1 | 7,762,396 | - | 6,681 | ND |
| *Streptomyces* sp. SirexAA E | ASM17719v2 | 7,414,440 | - | 6,663 | ND |
| *Streptomyces* sp. S8 | ASM209499v1 | 7,529,075 | 72,789 [1] | 6,624 [78] | 12,184 |
| *S. viridosporus* T7A ATCC 39115 | ASM870451v1 | 7,280,536 | - | 6,603 | ND |
| *S. luteoverticillatus* CGMCC 15060 | ASM397071v1 | 7,367,863 | - | 6,574 | ND |
| *S. pluripotens* MUSC 135 | ASM80224v2 | 7,346,075 | - | 6,561 | ND |
| *Streptomyces* sp. SAT1 | ASM165449v1 | 7,472,530 | - | 6,503 | ND |
| *S. ficellus* NRRL 8067 | ASM973990v1 | 7,078,240 | - | 6,412 | 21,921 |
| *S. koyangensis* VK A60T | ASM342892v1 | 7,220,839 | - | 6,365 | 20,855 |
| *S. tirandamycinicus* HNM0039 | ASM309751v1 | 7,289,495 | - | 6,363 | ND |
| *S. albidoflavus* SM254 | ASM157738v1 | 7,170,504 | - | 6,298 | ND |
| *S. ongiicola* HNM0071 | ASM312236v1 | 7,180,417 | - | 6,166 | ND |
| *S. fradiae* ATCC 10745 | ASM870442v1 | 6,725,579 | - | 5,877 | 48,424 |
| *S. seoulensis* KCTC 9819 | ASM432862v1 | 6,339,363 | 86,882 [1] | 5,866 [107] | 20,875 |
