## Supplementary_Table_3 for "Dynamics of the compartmentalized *Streptomyces* chromosome during metabolic differentiation"

### Table S3: Gene distribution in the central region *versus* the extremities (defined by the first and last rRNA operons) of *Streptomyces ambofaciens* ATCC23877 chromosome depending on features of interest

| Features | Number of genes of interest | % of genes of interest present within the extremities | % of genes within the central region presenting the feature of interest | % of genes within the extremities presenting the feature of interest | Odds ratio[a] | *p* value[b] |
|---|---|---|---|---|---|---|
| **Chromosomal genes** | 7128 | 46.5 | | | | |
| **Genome features related to the level of conservation** | | | | | | |
| **CDSs of the actinobacterial signature** | 72 | 16.7 | 1.6 | 0.4 | **0.23** | $1.4.10^{-7}$ |
| **Core CDSs** | 1020 | 11.3 | 23.7 | 3.5 | **0.12** | $<2.2.10^{-16}$ |
| **CDSs with a high level of persistence (>0.95)** | 2186 | 17.6 | 47.2 | 11.6 | **0.15** | $<2.2.10^{-16}$ |
| **GIs** | 1207 | 80.4 | 6.2 | 29.3 | **6.27** | $<2.2.10^{-16}$ |
| **Strain-specific CDSs[c]** | 158 | 85.4 | 0.6 | 4.1 | **6.99** | $<2.2.10^{-16}$ |
| **Genome features related to functional annotation** | | | | | | |
| **Functional RNA (excluding rRNA)** | 75 | 13.3 | 1.7 | 0.3 | **0.17** | $1.3.10^{-9}$ |
| **CDSs encoding functions involved in translation** | 175 | 13.7 | 4.0 | 0.7 | **0.18** | $<2.2.10^{-16}$ |
| **NAP genes** | 49 | 26.5 | 0.9 | 0.4 | **0.41** | 0.005798 |
| **SMBGCs** | 385 | 76.4 | 2.4 | 8.9 | **3.98** | $<2.2.10^{-16}$ |
| **Features related to the level of expression** | | | | | | |
| **Genes always expressed at very high level (CAT_4)[d]** | 332 | 11.7 | 7.7 | 1.2 | **0.14** | $<2.2.10^{-16}$ |
| **Genes always expressed at very low level (CAT_0)[d]** | 849 | 72.3 | 6.2 | 18.5 | **3.46** | $<2.2.10^{-16}$ |

[a]The odds ratio represents the odds than an outcome will occur within the extremities compared to the odds of the outcome occurring in the central region. Genes which distribution is statistically enriched in the central region or within the extremities are written in blue or in red, respectively. The genetic information of each TIR was taken into consideration for the calculations.

[b]The statistical significance of the odd ratios was assessed by a Fisher's exact test for count data.

[c]For this calculation, only the chromosomal strain-specific CDSs were taken into account (excluding the 44 strain-specific CDSs present on pSAM1 non-integrative plasmid).

[d]'always' refers to all the conditions investigated in this study (i.e. C1 to C7). Only chromosomal genes were considered in the calculation.
